# An incoherent feedforward loop interprets NFκB/RelA dynamics to determine TNF-induced necroptosis decisions

**DOI:** 10.1101/2020.04.30.070888

**Authors:** Marie Oliver Metzig, Ying Tang, Simon Mitchell, Brooks Taylor, Robert Foreman, Roy Wollman, Alexander Hoffmann

**Author notes:** Brighton and Sussex Medical School, University of Sussex, Brighton, BN1 9PX, United Kingdom (Simon Mitchell).

## Abstract

Balancing cell death is essential to maintain healthy tissue homeostasis and prevent disease. Tumor necrosis factor (TNF) not only activates nuclear factor κB (NFκB), which coordinates the cellular response to inflammation, but may also trigger necroptosis, a pro-inflammatory form of cell death. Whether TNF-induced NFκB cross-regulates TNF-induced necroptosis fate decisions is unclear. Live-cell microscopy and model-aided analysis of death kinetics identified a molecular circuit that interprets TNF-induced NFκB/RelA dynamics to control necroptosis decisions. Inducible expression of TNFAIP3/A20 forms an incoherent feedforward loop to interfere with the RIPK3-containing necrosome complex and protect a fraction of cells from transient, but not long-term TNF exposure. Furthermore, dysregulated NFκB dynamics often associated with disease diminish TNF-induced necroptosis. Our results suggest that TNF’s dual roles in either coordinating cellular responses to inflammation, or further amplifying inflammation are determined by a dynamic NFκB-A20-RIPK3 circuit, that could be targeted to treat inflammation and cancer.

## INTRODUCTION

The cytokine tumor necrosis factor (TNF) mediates diverse cell fate decisions in response to inflammation (Supplementary Figure 1A)^1, 2^. TNF-induced activation of nuclear factor κB (NFκB) regulates the expression of hundreds of inflammatory response genes involved in eliminating pathogens, resolving inflammation and healing^3, 4^. However, TNF is also a cell killing agent^2, 5^ and may trigger apoptotic or necroptotic cell death programs with distinct pathophysiological consequences^6^. While apoptotic cells fragment into membrane bound vesicles, which allows their removal and resolution of inflammation^7^, necroptotic cells spill damage-associated molecular patterns (DAMPs) into the microenvironment, which promotes inflammation^8, 9^. Indeed, necroptosis has been linked to acute and chronic inflammatory diseases^10–12^, and inhibition may be a promising therapeutic strategy^13^. Conversely, necroptosis may be beneficial in apoptosis-resistant cancer^14–17^ and to evoke an anti-tumor immune response^18–20^. However, too little is known about the regulatory network controlling necroptosis to allow for predictable manipulation as a therapeutic strategy^21^.

When an isogenic cell population is challenged with a cytotoxic stimulus, not all cells make the decision to die at the same time, and some cells may even survive altogether^22–26^. In principle, stimulus-induced cell fate decisions may merely be a function of the cell’s propensity to make that decision. For instance, TNF-related apoptosis-inducing ligand (TRAIL) sorts cells into survivors or non-survivors based on the state of the molecular signaling network^23^, and thus the fate decision of an individual cell is predictable prior to administering the stimulus^27^. Alternatively, the stimulus may trigger both promoters and inhibitors of the cell fate decision. In this case, the fate decision of an individual cell is determined by the competition of regulators, whose induction follows specific dynamics. The regulatory motif, known as an Incoherent Feedforward Loop, is thus known to have the capacity for differentiating the duration of the incoming stimulus^28^.

TNF’s cytotoxic activity was initially described in the L929 fibroblast cell line^5^ leading to the characterization of necroptotic cell death^29^. TNF is now known to first trigger the formation of signaling complex I by recruiting receptor interacting serine/threonine kinase 1 (RIPK1) to TNF receptor 1 (TNFR1), leading to the activation of the inhibitor κB kinase (IKK) and transcription factor NFκB^30^. Dissociation of RIPK1 from the plasma-membrane-bound complex I then allows for the recruitment of RIPK3 (complex IIb or the necrosome), which leads to RIPK3-oligomerization and phosphorylation of mixed lineage kinase like (pMLKL), causing plasma membrane rupture and necroptotic cell death^31–34^.

Prior studies investigated NFκB as a potential necroptosis inhibitor^35–38^, but in certain circumstances, NFκB may even promote necroptosis, e.g. by contributing to TNF production^15^. Furthermore, it remains unclear whether prior NFκB activity determines the propensity for cells to die, or whether TNF-induced NFκB activation may determine the decision making, which would require de-novo protein synthesis to be induced rapidly enough to affect the transition of signaling complexes I and II. Only the latter would constitute an incoherent feed forward loop capable of distinguishing stimulus dynamics.

Here, we sought to determine whether and how TNF-induced NFκB activation regulates TNF-induced necroptosis decisions. Given the potential of perturbation studies to skew the true regulatory network^39, 40^, we developed a live-cell microscopy workflow to study unperturbed L929 cells and obtain time-resolved quantitative necroptosis rates following TNF exposure. A conceptual mathematical model informed us how these death rate dynamics can be interpreted, leading us to identify TNF-induced and NFκB-responsive TNFAIP3/A20 as a key regulator of necroptotic fate decisions. The A20 circuit forms an incoherent feedforward loop to protect a fraction of cells from transient TNF doses, but renders them sensitive to long-term TNF exposure. As predicted by a more detailed mathematical model, dysregulated NFκB dynamics diminish the cell’s ability to make necroptosis decisions based on the duration of TNF exposure.

## RESULTS

### Necroptosis kinetics are reflective of an incoherent feedforward loop

To distinguish whether TNF-induced necroptosis decisions are merely a function of a preexisting propensity or the dynamics of stimulus-induced regulators, we constructed two simple conceptual models. In the first, TNF induces activation of RIPK1/3 and the necroptosis effector pMLKL, but this signaling is counteracted by an unknown, constitutively expressed survival factor (Figure 1A). In the second model, TNF also induces inhibitor of κB (IκB)-controlled NFκB^41^, which in turn induces expression of the survival factor, thus forming an incoherent feedforward loop (Figure 1B)^28, 42, 43^. To account for cell-to-cell heterogeneity, we assumed stochastic gene expression^44, 45^ of the survival factor, such that repeated simulations corresponded to different cells that have different steady state amounts, and applied an arbitrary threshold for pMLKL corresponding to irreversible cell death (Figure C and D; Supplementary Notes).

**Figure 1.**
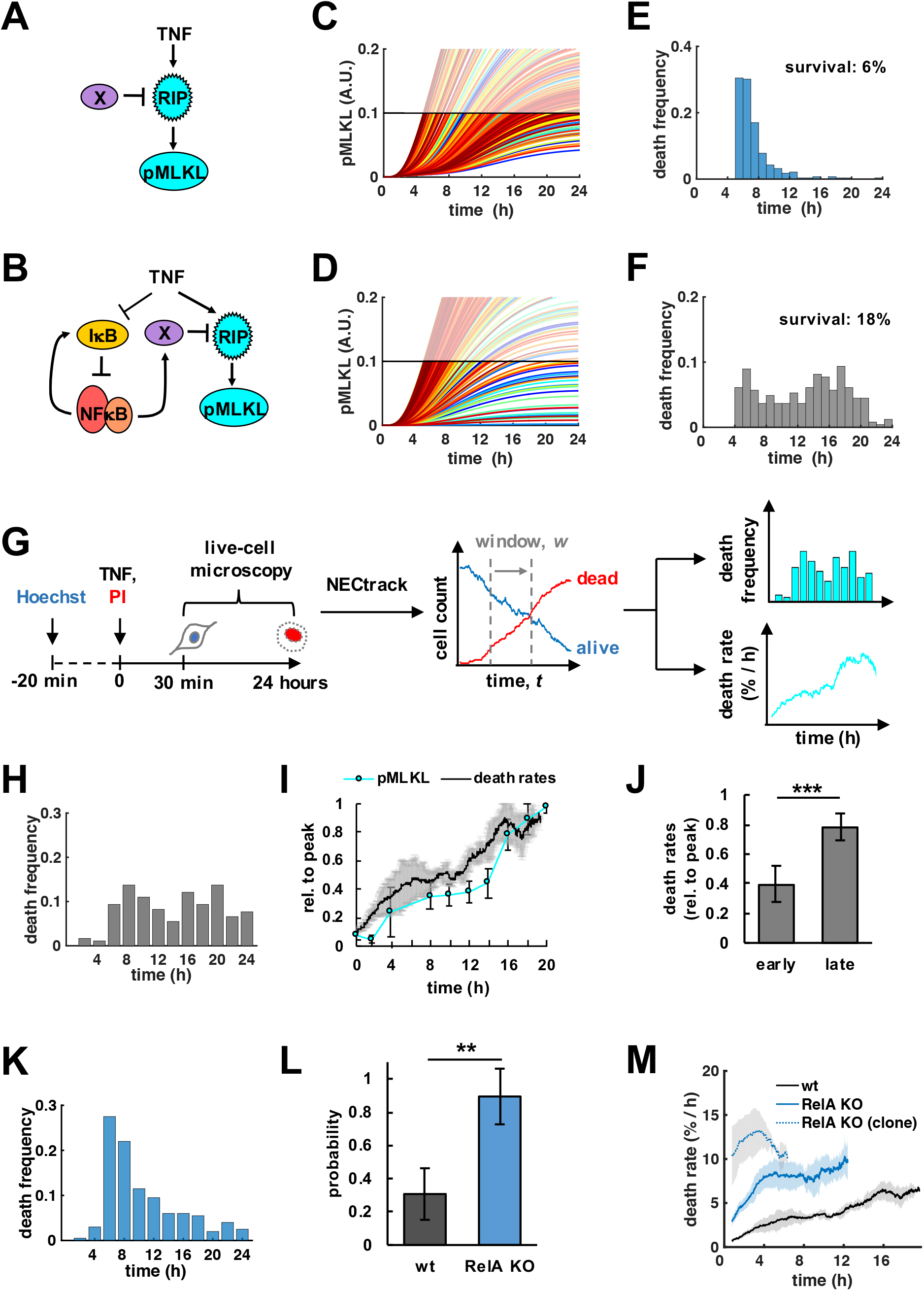
Necroptosis kinetics are reflective of an incoherent feedforward loop. (A, B) Conceptual mathematical modeling schematics depict TNF-induced necroptosis signaling via RIPK1/3 (RIP) and phosphorylation of MLKL (pMLKL). RIP is counteracted by a putative (A) constitutive, or (B) stimulus-induced, NFκB-dependent survival factor X. (C, D) Time course simulations of pMLKL levels in 300 single cells where each trajectory crossing a threshold represents a cell death event. (E, F) Distributions of death times that result from simulations in (C, D), respectively. Fractional survival indicated after 24-hour time course simulation. (G) Livecell microscopy workflow and automated image analysis via *NECtrack* to quantify TNF-induced necroptosis kinetics in L929 cells. Distributions of death times and death rates are computed from raw counts of live and dead cells based on nuclear propidium iodide (PI) staining. (H) Distribution of death times in TNF-treated L929 wildtype (wt) cells (representative data of three independent experiments). (I) Normalized death rates in L929 wt cells plotted with pMLKL protein levels measured via immunoblot (mean of three independent experiments ± standard deviation; corresponding images of representative Western blot experiment in Supplementary Figure 1E). (J) Average death rates of the early (<12 hours) and late phase of the TNF time course data in (I) (mean of three independent experiments ± two-tailed Student’s t-test ***P<0.001). (K) Distribution of death times in L929 RelA-knockout (RelA KO) cells treated with TNF (representative data of three independent experiments). (L) Probability of unimodal distributions of death times calculated by Hartigan’s dip significance (mean of three independent experiments ± standard deviation; two-sample t-test **P<0.01). (M) Death rates in TNF-treated cell lines including clonal RelA-knockout population (mean of three independent experiments ± standard deviation).

Plotting the cell death time course by hourly binning the number of simulations in which pMLKL exceeds the threshold, we found that when the survival factor is pre-existing and the mechanism constitutive, death times followed a unimodal distribution (Figure 1E). In contrast, TNF-induced, NFκB dependent expression of a pro-survival factor produced a bimodal death time distribution (Figure 1F). While exact death times are a function of the particular parameters chosen in this analysis, the distinction between unimodal *vs*. bimodal death time distributions was a robust feature of pre-existing *vs*. inducible survival mechanisms (Supplementary Notes).

Next, we established the live-cell microscopy workflow and automated image analysis tool *NECtrack* to measure TNF-induced necroptosis dynamics (Figure 1G, Supplementary Video). TNF-treated L929 cells were imaged and tracked for 24 hours by nuclear Hoechst staining, and new necroptotic death events were identified by nuclear uptake of propidium iodide (PI) added to the culture medium. This workflow quantified necroptosis without being confounded by concurrent cell proliferation (Supplementary Figure 1B), a common bias of bulk readout assays based on fractional survival^46^. Average necroptosis rates per hour were obtained by normalizing new death events to the number of present live cells. The use of nuclear dyes at low concentrations had no significant effect on cell numbers (Supplementary Figure 1C). To address the concern of phototoxicity, we compared different counting protocols and found that the microscopy workflow actually preserved cell viability better than a parallel, but independent counting protocol that required trypsinization (Supplementary Figure 1D).

Our measurements indicated a bimodal distribution of death times in L929 cell populations undergoing TNF-induced necroptosis (Figure 1H), reflected by two-phased death rate dynamics (Figure 1I). Indeed, death rates were about two-fold higher in the late vs. early (< 12 hours) phase (Figure 1J), and correlated with levels of pMLKL, the molecular marker for necroptosis (Figure 1I, Supplementary Figure 1E). Similar biphasic necroptosis kinetics were observed in a newly cloned L929 cell population treated with TNF (Supplementary Figure 1F). These experimental results suggested that TNF does not only trigger pro-death signaling leading to necroptosis, but also activates a mechanism that provides for transient protection of a fraction of cells within the population.

We asked whether TNF-induced activation of NFκB RelA (Supplementary Figure 1G and H) may be responsible. Interestingly, over a time-course of 24 hours, RelA activity and death dynamics were inversely correlated (Supplementary Figure 1I). Depriving L929 cells of RelA via CRISPR/Cas9 (Supplementary Figure 1J) led to similar fractional survival after 24 hours (Supplementary Figure 1K), but shifted necroptosis to the early phase resulting in a largely unimodal distribution of death times (P= 0.9, Figure 1K and L) and single-phased death rates (Figure 1M). Increased necroptotic cell death during the early phase was correlated with detection of pMLKL (Supplementary Figure 1L) with most cells displaying morphological characteristics of necrotic cell death as expected (Supplementary Figure 1M and N). This sensitizing effect was even more pronounced in a clonal knockout population (Figure 1M) that was confirmed to be fully RelA-deficient (Supplementary Figure 1J). Together, these results support that TNF-induced RelA is a potent inhibitor of necroptosis transiently protecting a fraction of L929 cells from necroptosis.

### Rapid induction of A20 transiently inhibits the RIPK1-RIPK3 complex and necroptosis

Next, we sought to identify the inducible mechanism by which RelA transiently protects a fraction of L929 cells from necroptosis. NFκB-induced gene products may limit necroptosis by inhibiting reactive oxygen species (ROS) production or pro-death c-Jun N-terminal kinase (JNK) signaling^37, 47–50^. While TNF-induced generation of ROS may amplify necroptosis signaling^51^, addition of the anti-oxidant butylated hydroxyanisole (BHA) (Supplementary Figure 2A), or the specific JNK inhibitor SP600125 (Supplementary Figure 2B) had limited effect in RelA-knockout cells. This suggested that RelA-mediated inhibition of necroptosis was not critically mediated by ROS or JNK.

Several potential NFκB-responsive target genes have been described to modulate TNFR-induced complex I, complex II and/or the necrosome^38, 52^. In complex IIa, FLIP-L has been implicated in restricting the proteolytic activity of pro-caspase-8 and directing its substrate specificity towards RIPK1 to disassemble the RIPK1-RIPK3-complex^53–56^. However, FLIP-L mRNA was not induced by TNF in L929 cells (Supplementary Figure 2C), and similar dynamics of FLIP-L cleavage were observed in wildtype and RelA-knockout cells (Supplementary Figure 2D), indicative of comparable proteolytic activity of pro-caspase 8 in complex II^54, 57^. Similarly, mRNA levels of CYLD or cIAP1, which are involved in the regulation of complex I activity^21^, were not significantly induced by TNF, or reduced in RelA-knockout cells (Figure 2A). In contrast, TNFAIP3/A20 and cIAP2 were significantly induced within 0.5 or 1 hours of TNF treatment in a RelA-dependent manner (Figure 2A), whereas only A20 expression was transient and correlated with dynamics of RelA activity (Supplementary Figure 2E). Inducible mRNA expression was accompanied by elevated A20 protein detected after 2 hours of TNF treatment in wildtype, but not RelA-knockout cells, while constitutive expression levels were not significantly affected (Supplementary Figure 2F). Thus, inducible A20 appeared to be a promising candidate to mediate RelA-dependent transient protection from necroptosis.

**Figure 2.**
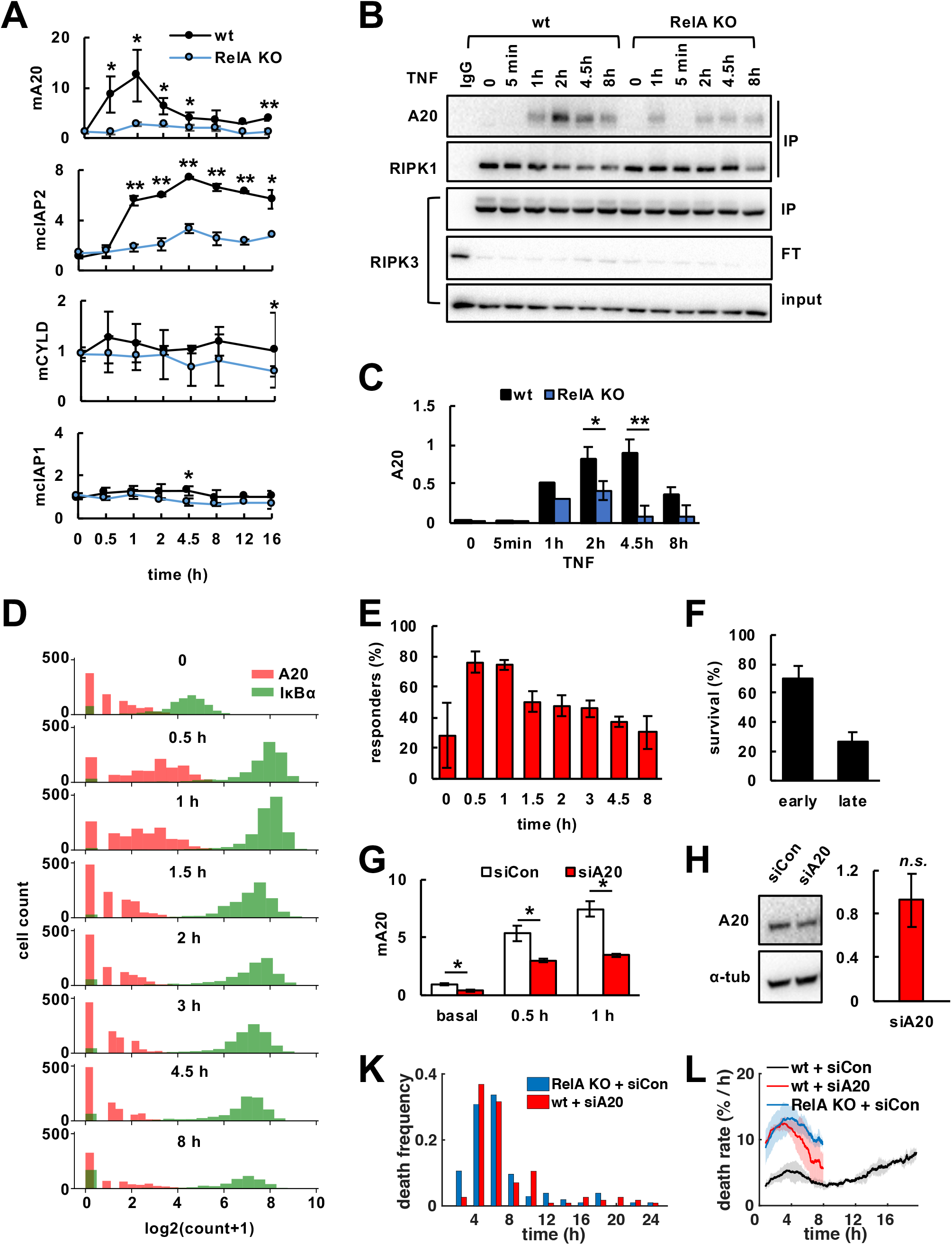
Rapid induction of A20 transiently inhibits the RIPK1-RIPK3 complex and necroptosis. (A) TNF-induced mRNA expression in indicated L929 cell lines measured via qRT-PCR (mean of three independent experiments ± standard deviation, two-tailed Student’s t-test *P<0.05, **P<0.01). (B) Immunoblot after co-immunoprecipitation (IP) of RIPK3. FT, flow through. (C) Relative quantification of A20 in RIPK3-IP fraction (means and statistical significance established for 2 and 4.5 hour time points from three independent experiments ± standard deviation; two-tailed Student’s t-test *P<0.05, **P<0.01). (D) Histogram of volume normalized mRNA copy numbers measured by smFISH of NFκB target genes A20 and IkBa in L929 wildtype (wt) cells treated with TNF (representative data of three independent experiments; two additional independent experiments in Supplementary Figure 4H). (E) Fractions of “responder” cells (A20 count > 1 per cell) (mean of all three independent experiments ± standard deviation). (F) Fractional survival of wt cells after the early (<12 hours) and late phase of TNF time course obtained by microscopy (mean of three independent experiments ± standard deviation). (G) A20 mRNA expression via qRT-PCR in TNF-treated wt cells transfected with targeting (siA20) or non-targeting (siCon) siRNA (mean of three independent experiments ± standard deviation; twotailed Student’s t-test *P<0.05). (J) Immunoblot and relative quantification of basal A20 protein after siRNA-knockdown normalized to non-targeting siRNA control (mean of three independent experiments ± standard deviation; two-tailed Student’s t-test revealed no statistically significant difference between targeting and non-targeting siRNA treatment, n.s., P>0.05). (K) TNF-induced distributions of death times (representative data of three independent experiments), and (L) death rates obtained by live-cell microscopy (mean of three independent experiments ± standard deviation).

Several A20-dependent mechanisms to limit TNF-induced cell death have previously been reported^58–62^. In complex I, A20 binds and stabilizes M1-ubiquitin chains, which may limit formation of death-inducing complex II^58, 59, 61, 62^. In addition, A20 integrates into the downstream necrosome and restricts RIPK3 activation to limit necroptosis^60^. Whereas complex I forms rapidly^63^, the necrosome is activated slower^38^, potentially allowing *de novo* expressed A20 to interfere with its activity. Indeed, we found that inducible expression of A20 coincided with its increased dynamic integration into RIPK3 immuno-precipitates at 2 and 4.5 hours in wildtype compared to RelA-knockout cells (Figure 2B and C). This was accompanied by decreased binding of RIPK1 in wildtype cells (Figure 2B), indicating destabilization of the necrosome^60^. Similar levels of phosphorylated RIPK1 (Ser166) (Supplementary Figure 2G) and cleaved FLIP-L (Supplementary Figure 2D) were detected in wildtype and RelA-knockout cells, indicating that complex I and II activities^54, 57, 64–66^ were less affected by inducible A20.

As biphasic necroptosis kinetics had indicated that the RelA-dependent survival mechanism only protected a fraction of cells from premature necroptosis, we asked whether cell-to-cell heterogeneity in TNF-induced A20 expression may be responsible. Analyzing TNF-induced gene expression in single cells via smFISH revealed that A20 mRNA was only upregulated in 76% and 75% of “responder” cells at 0.5 and 1 hours, respectively, whereas the NFκB-responsive target IκBα was induced in nearly all cells (Figure 2D, E, Supplementary Figure 2H). In fact, the fraction of “responder” cells coarsely correlated with fractional survival after the early phase of TNF treatment (Figure 2F).

To further confirm the functional requirement of inducible A20, we performed siRNA-mediated knockdown of A20. Knockdown conditions were optimized to significantly decrease TNF-induced expression of the A20 mRNA (Figure 2G), but – due to protein half-life – without having a significant effect on basal A20 protein levels present at the start of the TNF stimulation (Figure 2H). These conditions strongly sensitized L929 wildtype cells to TNF leading to the majority of cells dying during the early phase, which resulted in unimodal death time distributions (Figure 2K) and single-phased death rate dynamics comparable to RelA-knockout cells (Figure 2L). In contrast, loss of cIAP2 (Supplementary Figure 2I, J) had no significant effect on necroptosis rates (Supplementary Figure 2K), which was in line with previous findings^51^.

Together, our results indicated that following TNF-induced, RelA-responsive A20 expression in a subset of cells, A20 binds to and subsequently inhibits RIPK1-RIPK3 complexes, thereby transiently protecting these cells from necroptosis. However, as RelA activity and A20 expression subside, these cells may become sensitive again and undergo necroptosis in the later phase of the time course.

### The NFκB-A20-RIPK3 incoherent feedforward loop discriminates TNF stimulus dynamics

To further investigate the crosstalk between TNF-induced NFκB and necroptosis fate decisions, we constructed a mathematical model that integrates the diverse roles of A20 in NFκB activation and necroptosis control (Figure 3A)^58–62, 67–70^. The model consists of 41 species and 98 reactions (Supplementary Figure 3A, Supplementary Notes), and combines the previously published modules for TNFR-IKK and IκB-NFκB signaling^69, 71^ with a newly constructed necroptosis module depicting activation of RIPK1, RIPK3 and the effector pMLKL. We parameterized the model based on literature, and to fit our experimental measurements in L929 cells (Supplementary Notes). Simulation of 24 hours of TNF treatment accurately recapitulated measurements of A20 expression at the mRNA and protein level (Supplementary Figure 3B, C), NFκB activation (Supplementary Figure 3D), and necrosome activity (Supplementary Figure 3E-G)^72^. Accounting for cell-to-cell heterogeneity in A20 expression as measured in smFISH experiments, as well as in RIPK1 activation (Supplementary Notes), we tested whether the model also recapitulated necroptosis kinetics in TNF-treated L929 cell populations, and the NFκB-dependent regulation. Indeed, simulations of wildtype (Figure 3B) or RelA-knockout cell populations (Figure 3C) showed the distinctive bimodal vs. unimodal distributions of death times as well as the two- vs. single-phased death rates, respectively, that we had observed in experiments. These results provided quantitative support to the notion that TNF-induced, RelA-mediated expression of A20 provides transient protection from necroptosis.

**Figure 3.**
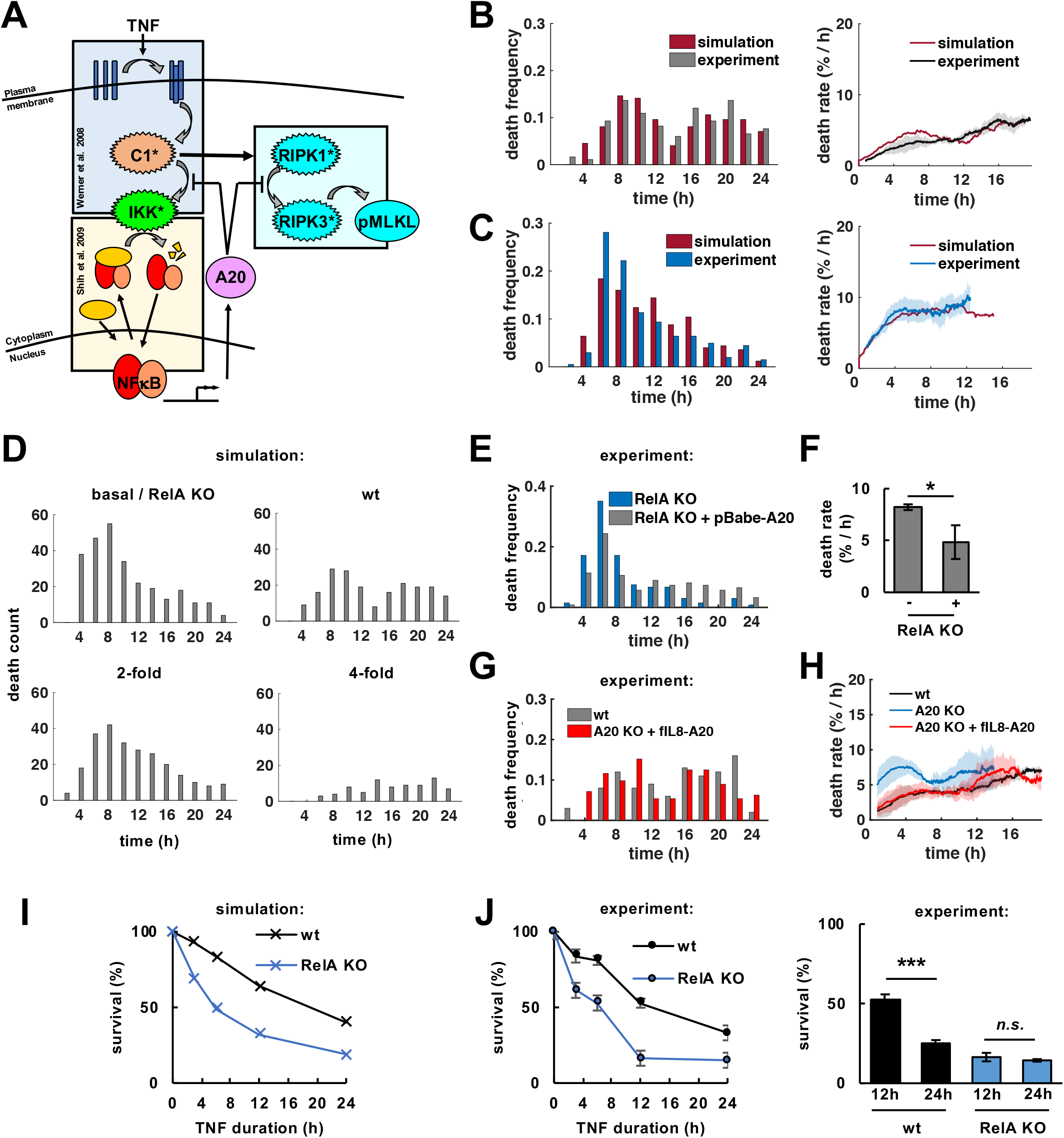
The NFκB-A20-RIPK3 incoherent feedforward loop discriminates TNF stimulus dynamics. (A) Modeling schematics depict TNF-induced activation of complex I (C1) and IKK to induce transcriptional activity of NFκB. C1 can also initiate activation of RIPK1 and RIPK3 to induce phosphorylation of necroptosis executor MLKL (pMLKL). TNF-induced expression of IκB attenuates NFκB, whereas A20 inhibits IKK and RIPK3. Computational simulations and microscopy analysis of death time distributions (left, representative data of three independent experiments) and death rates (right, mean of three independent experiments ± standard deviation) in TNF-treated L929 (B) wildtype (wt), or (C) RelA-knockout (KO) cells. (D) Simulated death time distributions with 2- or 4-fold increased constitutive A20 expression in the absence of inducible transcription (300 simulated cells per condition). (E) Death time distributions in TNF-treated parental RelA KO cells (-) or RelA KO cells expressing A20 from a constitutive transgene (pBabe-A20, +, representative data of three independent experiments). (F) Average death rates (<12 hours) in TNF-treated cells (mean of three independent experiments ± standard deviation; two-tailed Student’s t-test *P<0.05, or no statistically significant difference, n.s., P>0.05). (G) Distribution of death times (representative data of three independent experiments), or (H) death rates in TNF-treated wt, parental A20 KO cells, or A20 KO cells reconstituted with an NFκB-inducible transgene (fIL8-A20; mean of three independent experiments ± standard deviation). (I) Simulations and (J) experimental measurements of 24hour fractional survival after varying durations of transient or sustained (24h) TNF stimulation (mean of three independent experiments ± standard deviation; two-tailed Student’s t-test ***P<0.001, or no statistically significant difference, n.s., P>0.05).

Previous work demonstrated that while A20 plays a key role in modulating NFκB dynamics, its NFκB-inducible expression does not^69^. We wondered if the inducibility of A20 was instead required to provide proper dynamic regulation of TNF-induced cell death decisions. We employed the mathematical model, set the inducible expression of A20 to zero, and simulated 24 hours of TNF treatment with different levels of only constitutive A20 expression. While constitutively elevated A20 expression (2- or 4-fold) protected from death, two-phased necroptosis dynamics as characteristic in wildtype cells were not predicted (Figure 3D). To test this experimentally, we expressed A20 from a constitutive transgene in RelA-knockout cells, which led to 2.5-fold increased basal expression (Supplementary Figure 3H). Indeed, death dynamics remained unimodal (Figure 3E, Supplementary Figure 3I), although cells were protected from necroptosis compared to RelA-knockout cells (Figure 3F). However, when we reconstituted L929 A20-knockout cells (Supplementary Figure 3J) with an NFκB-inducible transgene (Supplementary Figure 3K), two-phased necroptosis dynamics were restored (Figure 3G and H, Supplementary Figure L). These results indicated that the TNF-inducible RelA-A20-RIPK3 circuit motif plays a critical role in shaping necroptotic death kinetics.

In physiological settings TNF is typically secreted in a transient manner, while pathologic conditions may be associated with prolonged TNF secretion. We hypothesized that the inducible RelA-A20-RIPK3 circuit may determine the fractional survival of cells in response to transient TNF stimulation, while leaving cells sensitive to long lasting TNF exposures. Model simulations testing a range of different temporal TNF doses (3-12 hours, Supplementary Notes) predicted that L929 wildtype cells would indeed be better protected from transient TNF exposures than RelA-knockout cells, while remaining sensitive when exposed to long-lasting TNF stimulation (Figure 3I). Experiments confirmed that wildtype, but not RelA-knockout populations, were able to discriminate short-term exposures of up to 12-hours from sustained 24-hour TNF treatment (Figure 3J). However, wildtype and RelA-knockout cells responded similarly to different TNF concentration doses (Supplementary Figure 5M), suggesting that the primary role of the RelA-A20-RIPK3 circuit motif is to discriminate between transient and sustained TNF, rather than concentration doses.

### Dysregulated NFκB dynamics diminish the cellular discrimination of TNF exposures

As dysregulated NFκB activity is often associated with disease^14, 73^, we utilized our mathematical model to explore necroptosis fate decisions as a function of altered RelA dynamics. To this end, we defined NFκB dynamics with an extrinsic pulse function rather than the normal IκB-circuit (Supplementary Notes). The model predicted that prolonged NFκB dynamics and A20 expression (Figure 4A) led to increased fractional survival in 24 hour-simulations of TNF treatment (Figure 4B). To experimentally test this scenario, we targeted the IκB regulatory system via CRISPR/Cas9-mediated gene knockout (Figure 4C), resulting in significantly prolonged TNF-induced RelA activity (Figure 4D, Supplementary Figure 4A, B), as well as prolonged expression of A20 mRNA and protein, while basal expression was unchanged (Figure 4E and F). As expected, IκBα/IκBε-knockout cells were more resistant to TNF-induced necroptosis with significantly increased fractional survival of 67% (Figure 4G) similar to the model prediction (56%, Figure 4B), and overall decreased death rates in response to 24 hours of sustained TNF treatment (Figure 4H). This effect was even more pronounced in a clonal population selected for IκBα/IκBε-knockout and CRISPR/Cas9-induced heterozygosity for p100 to compensate for upregulated IκB*δ* inhibitory activity^41^, while maintaining wildtype-like basal A20 expression (Figure 4H, Supplementary Figure 4C). Finally, siRNA-mediated knockdown targeting A20 in IκBα/IκBε-knockout cells confirmed that the protective effect was largely due to A20, as death rates now resembled those of wildtype cells treated with siA20 treatment (Figure 4I). Together, these data implicate that in conditions of dysregulated NFκB dynamics and prolonged expression of A20, cells are more likely to resist even long-lasting TNF exposures.

**Figure 4.**
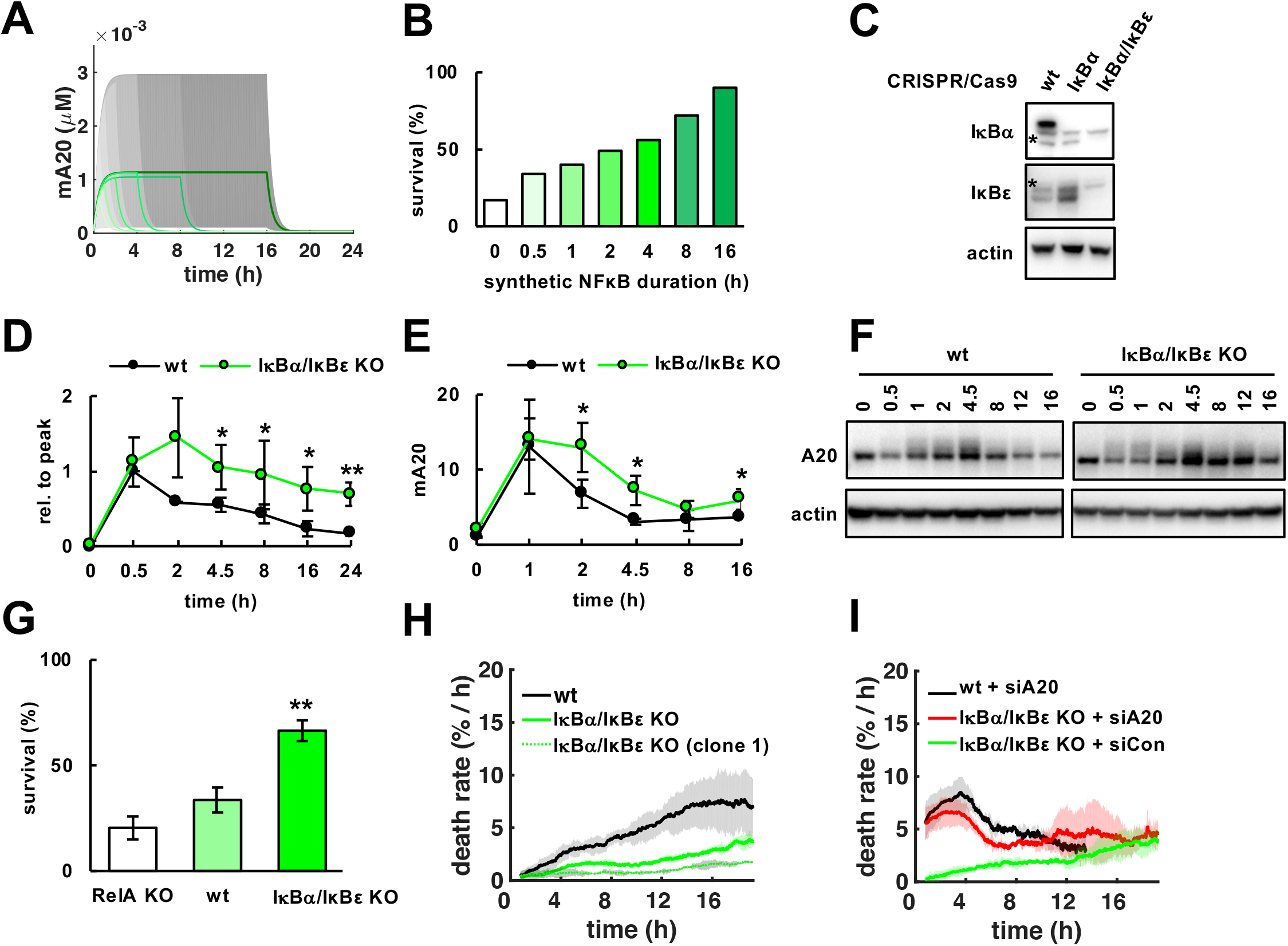
Dysregulated NFκB dynamics diminish the cellular discrimination of TNF exposures. (A) Simulations of A20 mRNA concentrations in versions of the NFκB-necroptosis model where expression is under the control of synthetic NFκB activity following step functions of 0.5, 1,2, 4, 8, or 16 hours duration (smoothed line is population average, and shaded area the 30^th^ percentile around the median). (B) Fractional survival that results from simulations in (A). (C) Immunoblot for IkBa and IkBε in L929 wildtype and CRISP/Cas9-knockout cell lines. Asterisks depict unspecific bands. (D) Normalized RelA activity dynamics after TNF treatment quantified via EMSA (mean of three independent experiments ± standard deviation; two-tailed Student’s t-test *P<0.05, **P<0.01; corresponding images of representative experiment in Supplementary Figure 4A). (E) A20 mRNA quantified via qPCR (mean of three independent experiments ± standard deviation; two-tailed Student’s t-test *P<0.05). (F) Immunoblot for A20 (representative data of three independent experiments). (G) Fractional survival after 24 hours of TNF treatment in indicated cell lines (mean of three independent experiments ± standard deviation; two-tailed Student’s t-test **P<0.01). (H) Death rates in TNF-treated indicated cell lines including isogenic IkBα/IkBε-knockout population (mean of three independent experiments ± standard deviation). (I) Death rates in TNF-treated cell lines treated with targeting (siA20) or non-targeting (siCon) siRNA (mean of three independent experiments ± standard deviation).

## DISCUSSION

In this study, we have addressed the regulatory mechanisms that determine TNF’s dual roles in inflammation, namely whether TNF elicits a cellular response that includes coordination and resolution of the inflammatory condition, or necroptotic cell death that may further amplify inflammation. Using time-lapse microscopy, we identified an incoherent feedforward loop involving TNF-induced NFκB/RelA activity and *de novo* expressed A20 protein, which provides potent, though transient protection to RIPK3-mediated necroptosis. We demonstrated that this molecular circuit ensures that a majority of cells survives transient TNF exposures, but, because of the transience of A20 expression, does not protect from long-lasting TNF exposure.

While a potential role of NFκB in inhibiting necroptosis was previously suggested^35–37^, the molecular regulatory circuits and its significance for necroptosis decision-making remained unknown. Although the anti-inflammatory protein A20 is a prominent NFκB-response gene, its robust TNF-inducibility is not required for inhibiting NFκB^69^, prompting the question of why A20 expression is so highly TNF-inducible. Here, we demonstrate that TNF-inducible A20 is in fact key to linking NFκB and the regulation of necroptosis decisions. Even under conditions of exceptional TNF-sensitivity as demonstrated in the L929 cell model system, NFκB-responsive A20 provides potent, though transient protection from necroptosis, which is critically determined by the duration of A20 expression and TNF exposure times. The A20 expression time course is controlled by NFκB dynamics, which is in turn a function of stimulus duration and IkB feedback regulation^69, 74^. We found that when cells are deprived of negative feedback mechanisms that ensure physiological NFκB dynamics, subsequent prolonged expression of A20 will diminish TNF-induced necroptosis.

Previous studies established A20 as an inhibitor of TNF-induced cell death^58–62, 68^. Via its ubiquitin binding domain ZnF7, A20 is believed to stabilize M1-linked ubiquitin chains in TNFR1-induced complex I, which may restrict complex II formation and thereby apoptosis and/or necroptosis as shown in MEFs^58^, macrophages^59^, and intestinal epithelial cells^61^. In addition, previous work in T cells and MEFs suggested that A20 binds to the necrosome, which may also be mediated via its ZnF7 domain, to enable ubiquitin editing and disruption of RIPK1-RIPK3-complexes^52, 60^. Our iterative approach of mathematical modeling and experiments provides a refined, quantitative and dynamic picture of A20’s roles in determining TNF-mediated fate decisions. While our results support that the amount of constitutively expressed A20 determines the overall propensity of cells to die, we show that inducible A20 expression kinetics critically shape the dynamics of TNF-induced necroptosis decisions. As upstream complex I is activated rapidly within minutes of TNF stimulation^63^, we reason that constitutively expressed A20 may integrate and limit the rate of transitioning into death-inducing complex II^75^ to determine both the apoptotic and necroptotic propensity. The active necrosome, however, forms within hours^38, 51^ and is therefore more susceptible to inducible A20 expression, which is why NFκB activity dynamics and the duration of the stimulus may be more critical in shaping necroptosis decisions. Indeed, our biochemical analyses showed that A20 expression kinetics coincided with its dynamic integration into RIPK1-RIPK3-complexes. Further systematic work will address how the distinct molecular mechanisms described for A20 in complex I/II and the necrosome quantitatively contribute to necroptotic and apoptotic death decisions.

Incoherent feedforward loops within biological signaling networks have previously been described to allow cells to dynamically adapt their gene expression^76^ and fate decisions in response to inflammation and p53-activating damage^77, 78^. Genetic and pharmacologic perturbation studies revealed that NFκB protects cells from apoptosis^63, 79^ by inducing the expression of anti-apoptotic target genes such as caspase inhibitor cFLIP^80^. In contrast to our findings about NFκB-mediated necroptosis control, however, TNF-induced apoptosis decisions may not be regulated by TNF-induced gene expression^63, 79^. While protein synthesis inhibitors sensitize cells to TNF-induced apoptotosis^80^, they also block constitutive protein expression, including key anti-apoptotic target genes such as cFLIP, whose short half-life requires continuous constitutive synthesis^39, 40^. Of note, cFLIP is only weakly induced by TNF ^40, 80^. The fact that TNF pulses as short as 30 seconds may be as effective as continuous exposure in eliciting apoptotic responses^81^ may suggest that the stimulus itself merely sorts cells by a preexisting apoptotic propensity, which may in turn be affected by the level of tonic NFκB activity^27, 82, 83^.

What might be the physiological consequences of the differential regulatory strategies by which NFκB controls apoptosis and necroptosis? A cell’s decision to undergo apoptosis appears to be inherent, depending on the general health of a cell, tonic NFκB, and hence its history of having responded appropriately to prior inflammatory conditions. If cells are unhealthy, they will be weeded out via apoptosis without causing much inflammation. In contrast, whether cells that express the necroptosis machinery will die of necroptosis is also a function of the dynamics of NFκB and duration of the TNF signal (Supplementary Figure 5A). Healthy NFκB activity dynamics in response to physiological TNF doses will ensure these cells participate in immune modulatory tissue processes rather than die. However, if TNF doses last longer, as they may in persistent infections, sepsis or chronic inflammatory diseases, cells may die via necroptosis and thereby release DAMPs to fuel an overwhelming inflammatory response (Supplementary Figure 7A). Indeed, loss of Caspase-8^84^ or FADD^85^ induces TNF-mediated necroptosis and inflammatory lesions in murine intestinal epithelium, resembling the pathology of inflammatory bowel disease. In turn, loss of MLKL or RIPK3 protected mice from TNF-induced systemic inflammatory response syndrome (SIRS)^10^. Two recent reports further pointed out that A20’s anti-inflammatory properties are not solely reliant on inhibiting IKK/NFκB, but depend on the prevention of TNF-induced cell death^59, 61^. In this context, our work establishes physiological NFκB dynamics as a safeguard against overwhelming inflammation, namely by securing physiological expression of A20 and therefore protecting from TNF-induced necroptosis.

In contrast, in tumors amplifying inflammatory responses via necroptotic cell death may have beneficial effects, increasing immunogenicity and helping to establish effective anti-tumor immunity (Supplementary Figure 5B). In this context, sensitizing cells to TNF or other necroptotic stimuli by counteracting inducible NFκB or the protective functions of A20 may have potential therapeutic value to enhance anti-tumor immunity.

## AUTHOR CONTRIBUTIONS

M.O.M. initiated, and M.O.M. and A.H. designed the study. M.O.M. performed the experiments and analyzed the data, with assistance from R.F. for smFISH experiments. M.O.M., S.M. and B.T. developed the automated image analysis tools for NECTrack. Y.T. performed the mathematical modeling. M.O.M. and A.H. interpreted the data and wrote the paper with valuable contributions by R.W.

## ACKNOWLEDGEMENTS

We are grateful to Tracy Johnson, Shubhamoy Ghosh and Sho Ohta for providing IκB-targeting CRISPR constructs. We thank Zhang Cheng for preparatory work on the NFκB module for the mathematical model, and Carlos Lopez for critical reading of the manuscript. This work was supported by NIH grants R01AI127867 and R01AI132731 to A.H., and with a Research Fellowship to M.O.M by the Deutsche Forschungsgemeinschaft (DFG).

