## Supplementary material for "An incoherent feedforward loop interprets NFκB/RelA dynamics to determine TNF-induced necroptosis decisions": Methods

### Materials

Mouse recombinant TNF (410-MT-10, R&D), propidium iodide (PI, P4864, Sigma Aldrich), Hoechst 33342 (H21492, Thermo Fisher Scientific), ZVAD-fmk (BML-P416-0001, Enzo Life Sciences), Necrostatin-1 (BML-AP309, Enzo Life Sciences), ROS inhibitor butylated hydroxyanisole (BHA, B1253, Sigma Aldrich), JNK inhibitor SP600125 (S5567, Sigma Aldrich).

### Cell culture

L929 cells were purchased from ATCC (NCTC clone 929 [L cell, L-929, derivative of Strain L] CCL-1) and maintained in DMEM (Corning) containing 10% FBS (Omega Scientific), 1% L-Glutamine and 1% penicillin/streptomycin (Thermo Fisher Scientific) at 5% $CO_2$ and 37°C. Isogenic L929 wildtype or CRISPR/Cas9-modified cell lines (RelA- or IκBα/IκBε-knockout) were established by single-cell sorting and clonal expansion as indicated.

### Imaging

Cells ($3 \times 10^5$) were seeded into eight-well µ-slides (ibidi) and grown for 24 hours. Following Hoechst staining (15 ng/ml for 20 min), propidium iodide (1 µg/ml), and TNF (10 ng/ml) were added to culture medium. After equilibrating for 30 minutes at 5% $CO_2$ and 37°C, cells were imaged using a Zeiss AxioObserver equipped with a 20X objective, LED (light-emitting diode) fluorescence excitation, and CoolSnap HQ2 camera. Differential interference contrast (DIC), Hoechst (excitation at 541 nm, cube BP515-560/FT580/LP590), and PI (excitation at 541 nm, cube BP515-560/FT580/LP590) images were taken every 1.5 minutes for 24 hours, and exported to MATLAB for automated analysis.

### Image analysis

Our automated image analysis tool NECtrack identifies, segments and tracks individual cells based on DIC and Hoechst images (Selimkhanov et al., 2014), and measures their mean nuclear PI intensity over time. Cells were declared dead when numerical threshold of nuclear PI was crossed for at least six consecutive time frames, and first frame was stored to generate histogram of death times. The status of each cell per frame was then integrated into a single binary matrix with cells and time points in rows and columns, respectively, with 0 representing "alive" and 1 indicating death had occurred. Absolute numbers of death events were converted to a rate of cell death proportional to the number of cells alive at a time point by using a five-hour sliding window, in which the number of new death events within a window was divided by the number of cells alive at the beginning of the window. The death rate at a time point is therefore the rate of death that will occur over the following five hours for cells alive at that time point; this accounts for continuous numerical changes in the population, e.g. by cell death or division. Average death rates per hour, i.e. the probability for individual cells to die within a given time window, is reported as the mean percentage of three independent experiments +/- standard deviations until remaining alive cell population drops under one third of the starting population (approx. 250-300 cells per experiment and condition). To obtain proliferative index, cell divisions were manually counted per 4-hour time window, and normalized to the number of alive cells present at the beginning of each window.

**Statistical analysis of distributions of death times**

All death times were placed into a single ordered array and passed to MATLAB's 'histcounts' function with 'Normalization' set to 'pdf' and histogram bins defined every 2 hours from 30 minutes (first time point) to 24.5 hours (final time point). The calculated probability distribution of cell death was plotted as a histogram. The null hypothesis that the sorted array of death times was drawn from a unimodal distribution was tested by Hartigan's dip significance test for unimodality, which calculates a probability of unimodality (Hartigan et al., 1985). This was repeated for three independent experiments for TNF treated wildtype or RelA-knockout cells. The probabilities of unimodality in each condition were compared with a two-sample t-test.

**Cell viability endpoint assays**

$1x10^4$ cells per well were grown in 96-well plates for 24 hours, and subjected to drug treatments. At respective endpoints, cells were stained with crystal violet, absorption was measured using a microplate reader (Epoch, BioTek), and normalized to mock-treated controls. As indicated, cells were pre-treated with ZVAD (30 μM), BHA (50 μM), or JNK inhibitor (10 μM) for 1 hour prior to TNF (10 ng/ml) treatment. In case of pulse stimulation experiments, cells were treated with TNF for 12 hours, washed and then incubated in regular medium before 24-hour endpoint measurement was taken. Alternatively, $6x10^5$ cells per well were grown in 6-well plates and treated with TNF for indicated durations, followed by trypsinization and manual counting of alive cells using a hemocytometer and Trypan blue exclusion staining.

**Co-immunoprecipitation and immunoblotting**

For Western blot analysis, 2-3x $10^6$ cells were grown in 10 cm plates for 24 hours. After treatment, dead cells were removed by thorough washing with PBS, and lysates of remaining adherent cells were prepared using RIPA buffer containing 1% Triton X-100 supplemented with PMSF, DTT and phosphatase inhibitors. Samples were normalized for protein amounts using a Bradford assay (BioRad). Detergent insoluble fractions were boiled for 10 min in 3X SDS sample buffer and directly subjected to gel electrophoresis. We used primary antibodies against pMLKL (ab196436, Abcam), RIPK1 (610459, BD Biosciences), pRIPK1 (31122S, Cell Signaling Technology), RIPK3 (PRS2283, Sigma Aldrich), RelA (sc-372, Santa Cruz), IkBα (sc-371, Santa Cruz), IkBβ (sc-945, Santa Cruz), IkBε (sc-7156, Santa Cruz), p52/100 (NR-145, generous gift from Nancy Rice), A20 (sc-166692, Santa Cruz), cFLIP (XA-1008, ProSci), Pan-cIAP (MAB3400, R&D), cIAP1 (ALX-803-335, Enzo Life Sciences), and corresponding secondary antibodies (HRP-conjugated, Cell Signaling). Signal was developed using chemiluminescent substrate (SuperSignal West Pico Plus, Thermo Fisher Scientific), visualized and quantified using ChemiDoc MP imaging system (BioRad). For co-immunoprecipitation experiments, 6x $10^6$ cells were grown for 24 hours in 15 cm plates and lysed in 30 mM Tris-HCL pH 7.4, 150 mM NaCl, 10% glycerol, 2 mM EDTA, 0.5% Triton, 0.5% NP-40, 1 mM DTT containing de-ubiquitinase inhibitor PR-619, protease and phosphatase inhibitors. Lysates were incubated with anti-RIPK3 for 4 hours, and protein G beads (Dynabeads, Thermo Fisher Scientific) for 1 hour to isolate complexes, followed by extensive washing in lysis buffer. If indicated, flow through was subjected to secondary co-immunoprecipitation with anti-RIPK1.

**Electrophoretic mobility shift assay (EMSA)**

Gel-shift assays were performed as previously described (Basak et al., 2007, Schroefelbauer et al., 2012). Briefly, nuclear extracts were incubated with $^{32}$P-labeled, double-stranded DNA probes containing kB binding sites in the presence or absence of anti-RelA or anti-RelB (sc-226, Santa Cruz), prior to nondenaturing acrylamide gel electrophoresis. Bands were visualized by autoradiography and quantified using ImageQuant software.

**Quantitative real-time PCR (qRT-PCR)**

RNA was purified using Direct-zol RNA Miniprep Plus Kit (Zymogen), and cDNA synthesized with iScript cDNA Synthesis Kit (Bio-Rad). qRT-PCR was performed with SYBR Green PCR Master Mix reagent using the D(DCt) method with RPL as normalization control, relative to unstimulated and stimulated signals in L929 cells to derive fold induction (A20: 5' – CCAGAGATTCCATGAAGCAAGA – 3', and 5' – CACATTTCAGCCTTGAGGCTAC – 3', cIAP1:

5’ – GAAAATGCTGACCCTACAGAGAC – 3’, and 5’ – CACCAGGCTCCTACTGAAGC -3’, cIAP2: 5’ – GAATGCAGACGCAGCAATC – 3’, and 5’ – CACCAGGCTCCTACTGAAGC -3’, CYLD: 5’ – AGCGTGACACAGGAAAGGAG – 3’, and 5’ – CCATGGATTTTTGGACTTGG – 3’, FLIP-L: 5’ – CCTCCAGCTCATCCTCTGTG – 3’, and 5’ TTTGTCCATGAGTTCAACGTG – 3’).

**Single-molecule fluorescence in situ hybridization (smFISH)**

Oligos (50 bp) were designed using custom software developed by the Zhuang lab for MERFISH (Moffitt et al., 2016). These oligos were comprised of a 30-bp region complementary to A20 or IκBα, and a 20-bp mouse orthogonal sequence that binds to dye labeled ‘readout oligo’. 52 different gene targeting regions were selected to tile along the length of the transcripts of interest. Dye-labeled oligos used Cy5 and Atto 565, respectively, to image labeled transcripts as diffraction limited spots. Cells (3 x $10^5$) plated in eight-well ibidi dishes, grown for 24 hours, and stimulated with 10 ng/ml TNF for different durations were fixed with 4% paraformaldehyde in PBS buffer. After rinsing with PBS, cells were permeabilized with 0.5% v/v Triton X-100 in PBS, followed by 3X rinse with Tris-buffered 300mM NaCl supplemented with 0.1% tween-20 (TB2XStw). Following equilibration for 10 minutes in TBS2Xtw supplemented with 30% formamide (MW), liquid was aspirated, and TBS2Xtw supplemented with 30% formamide, 10% dextran sulfate was added. After hybridizing with 50nM oligo mixture for both A20 and IκBα overnight (16 hours) at 37°C, cells were washed 2X with MW at 47°C for 30 minutes, and hybridized with a PER amplifier oligo at 25nM concentration in MH for 30 minutes at room temperature (https://www.nature.com/articles/s41592-019-0404-0). Cells were washed, stained for nuclei with DAPI, and stained with readout oligo in TB2XStw supplemented with 10% ethylene carbonate for 30 minutes, before being washed with the same solution without readout oligo. Imaging buffer (4mM PCA and 0.3U/mL of rPCO oxygen scavenging system in TBS2Xtw supplemented with 0.1 v/v murine RNAase inhibitor) was added before imaging on a custom configured Zeiss Axio Observer Z1 with 63X planapo objective, Zyla 4.2 sCMOS, and custom LED light engine built for MERFISH (Foreman et al., 2019). Images were background subtracted by difference of gaussian filtering with a high pass filter of 2.5 pixel sigma and 0.8 pixel low pass blurring filter. Local maxima were detected, and a 3x3 pixel region around local maxima intensity averaged. Number of spots found as a function of spot intensity was used to hysteresis threshold background spots from mRNA spots similar to Tsanov et al., 2016. Nuclei were segmented using watershed seed with smoothed local maxima in valleys of the negative smoothed intensity of the DAPI channel. Cells were imaged in a Green channel that produces smooth autofluorescence around the cell cytoplasm, and cytoplasm was segmented using the nuclei segmentation as seeds in a valley of negative autofluorescence intensity. RNA spots were assigned to cells if the cellular segmentation mask contained a particular spot. Cellular areas were calculated for each cell and used in volume normalization of RNA counts. Raw counts were normalized by dividing the count by the area of each cell, and multiplying by the average area of all cells to rescale counts back to the same means as before normalization. These volume normalized counts were log2 transformed with a pseudo-count of 1. Fraction of responders was calculated by finding the fraction of cells with an A20 count > 1 transcript per cell at each timepoint.

**CRISPR/Cas9-gene editing**

Guide RNAs (gRNAs) targeting RelA (5' – CACCG AATCGCATGCCCCGTTGCTT – 3', and 5' – CACCG TGTTCGATGATCTCCACATA – 3'), cIAP1 (5' – CACCG TGAATGCCACCTCGTTCCAG – 3', and 5' – CACCG TGGTCATCTAGTAGTCTGCC – 3'), cIAP2 (5' – CACCG ATGGTGCTCATCGCCGTGGA – 3', and 5' – CACCG TAAAGTGTGTATGGACCGAG – 3'), A20 (5' – CACCG TTTGCTACGACACTCGGAAC – 3', and 5' – CACCG CTCGGAACTTTAAATTCCGC – 3'), IkBα (5' – CACCG GGGTGCTGATGTCAACGCTC – 3', and 5' – CACCG CTGCGTCAAGACTGCTACAC – 3'), IkBε (5' – CACCGGCAACAGAATAGCACCGACG – 3', and 5' – CACCGAGCAACAGAATAGCACCGAC – 3'), or p100 (5' – GCCCGAGCGTTGCTGGACTA– 3', and 5' – CGCCGTAGTCCAGCAACGCT – 3') were cloned into lentiCRISPR v2 (52961, Addgene) (PMCID: PMC4486245) and used for lentivirus production in HEK293T cells. Infected L929 cells were selected with Puromycin (8 ug/ml) until cell death subsided, and subjected to single-cell sorting and clonal expansion as indicated. Knockout was confirmed by Western blot or – in the case of cIAP2 – by HRM analysis. To this end, genomic DNA was isolated using Quick-DNA Miniprep Plus (Zymogen), subjected to PCR amplification (5' – ACAGTCCCATGGAGAAGCAC – 3', and 5' – CTTGTGCTCAAAGCAGGACA – 3'), and subsequent melt curve analysis using SYBR Green (BioRad) and temperature increments (0.2°C steps).

**Transfection of short interference (si) RNA**

Reverse transfection of L929 cells was performed using transfection reagent (DharmaFECT1) and siRNAs (non-targeting control: D-001206-13, A20: M-058907, Dharmacon) at a final concentration of 50 nM. Knockdown efficiencies were tested prior to and in response to TNF treatment on mRNA and protein level as indicated.

**Expression of A20 transgenes**

pBabe-A20 containing a human A20 ORF was used for retroviral production (Werner et al., 2008) and infection of L929 RelA-knockout cells. In L929 A20-knockout cells, inducible A20 was reconstituted from transgene fIL8-A20 containing A20 ORF under the control of the IL8-promoter (Lois et al., 2002, Werner et al., 2008).

**Data reproducibility and statistical analysis**

Each experiment was repeated at least three independent times and performed in different weeks. All measurements were taken from distinct cellular samples, rather than measuring the same sample repeatedly. Microscopy analysis and all other data or images presented in the same figure panel were taken from measuring the respective conditions side-by-side in the same independent experiment. Statistical analysis was performed on means of these three individual experiments using two-tailed Student's t-test with P values noted in respective figure legends.

**Data availability**

Source data including uncropped images (microscopy, immunoblots) and microscopy analysis data are available from the corresponding author upon request. The Supplementary Video is in the repository Dryad with the download link:
https://datadryad.org/stash/share/Qgk61aoXPwWCLL7u-GNNOR6fD4F9aTdogkUF-si8_v8.

**Code availability**

The NECtrack package (including sample datasets) and NFκB-necroptosis modeling code (including all source data necessary to generate plots displayed in this study) are in the repository Dryad with the download link:
https://datadryad.org/stash/share/Qgk61aoXPwWCLL7u-GNNOR6fD4F9aTdogkUF-si8_v8.
