## Supplementary Figures for "An incoherent feedforward loop interprets NFκB/RelA dynamics to determine TNF-induced necroptosis decisions"

### Supplementary Figure 1

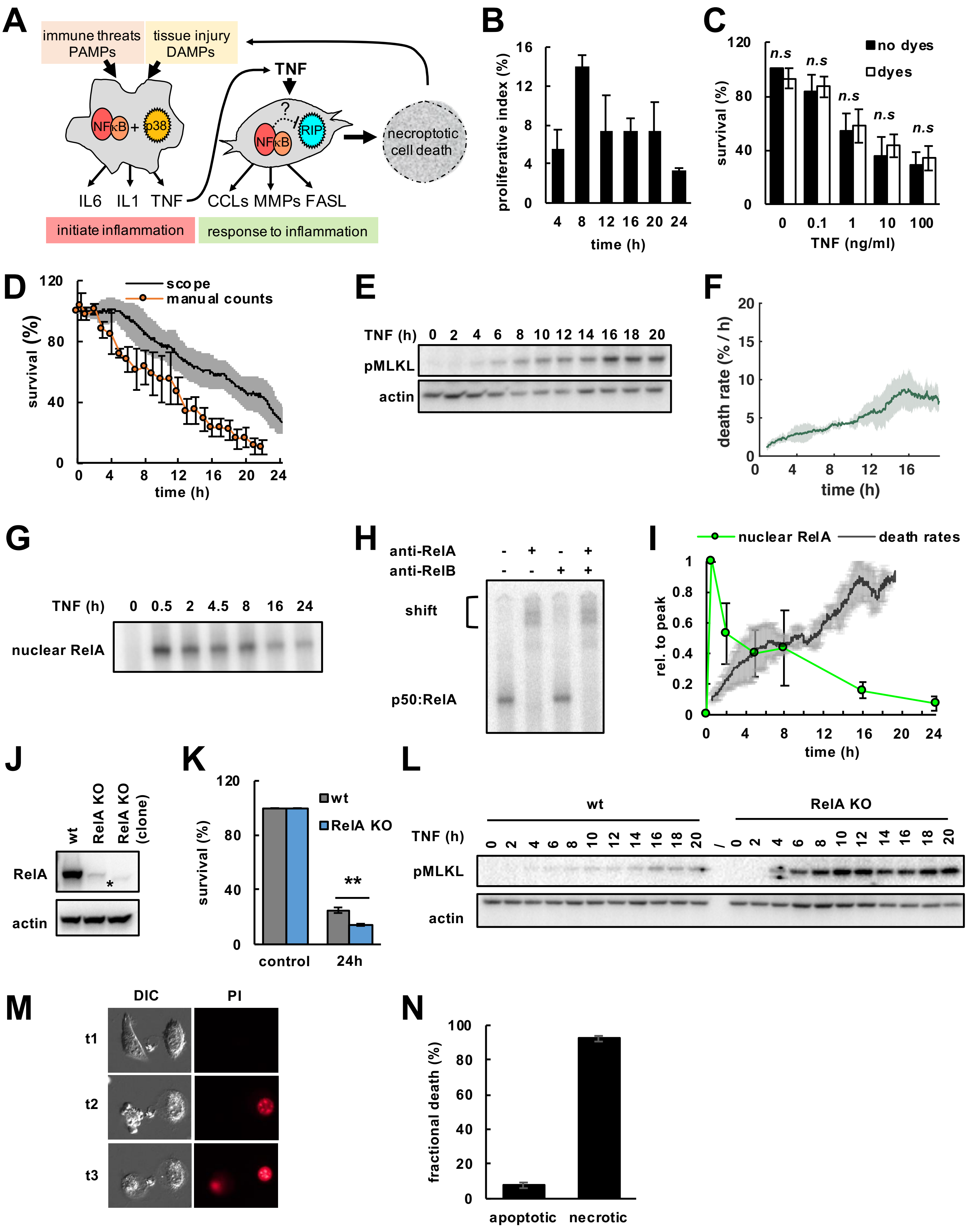

### Supplementary Figure 2

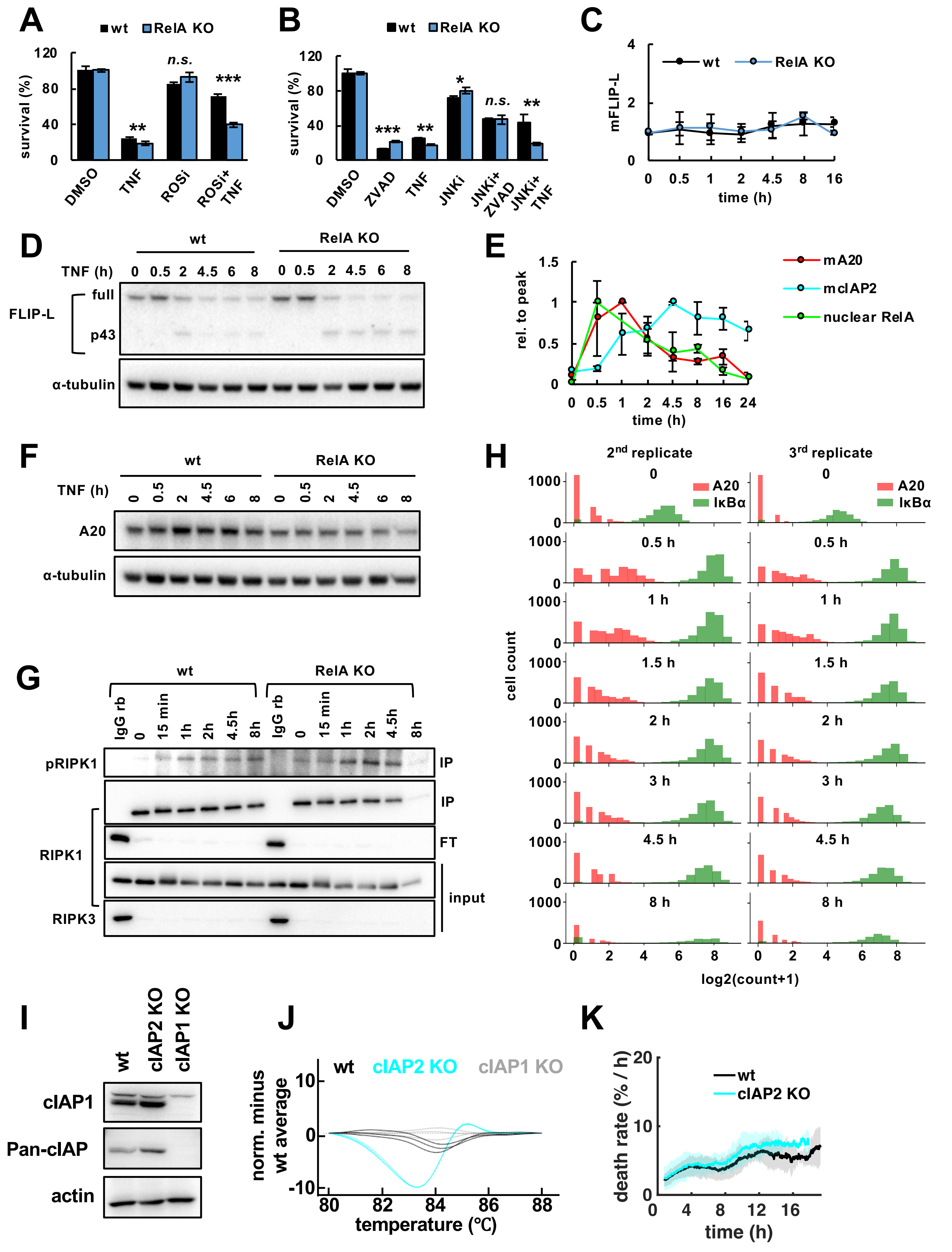

### Supplementary Figure 3

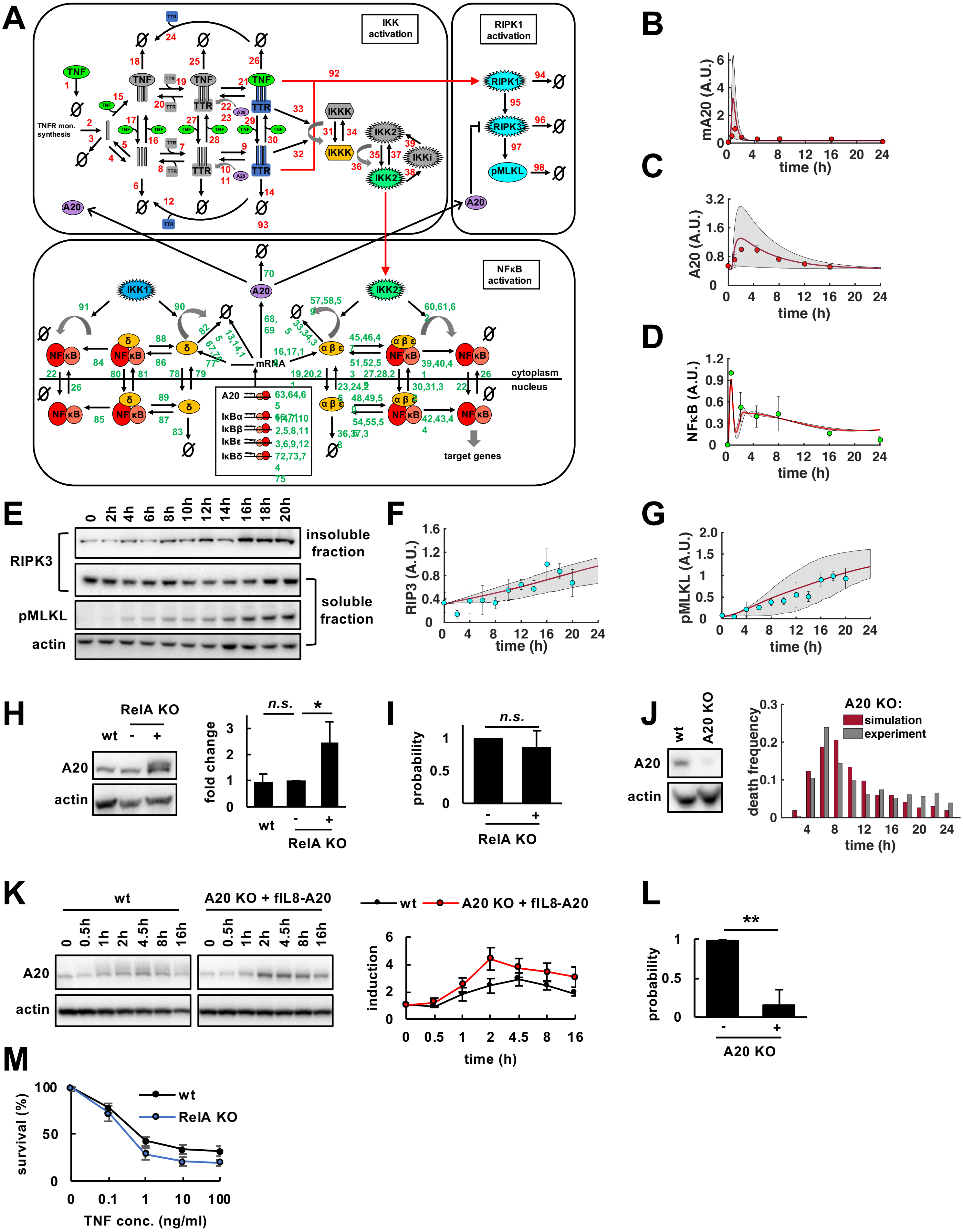

### Supplemental Figure 4

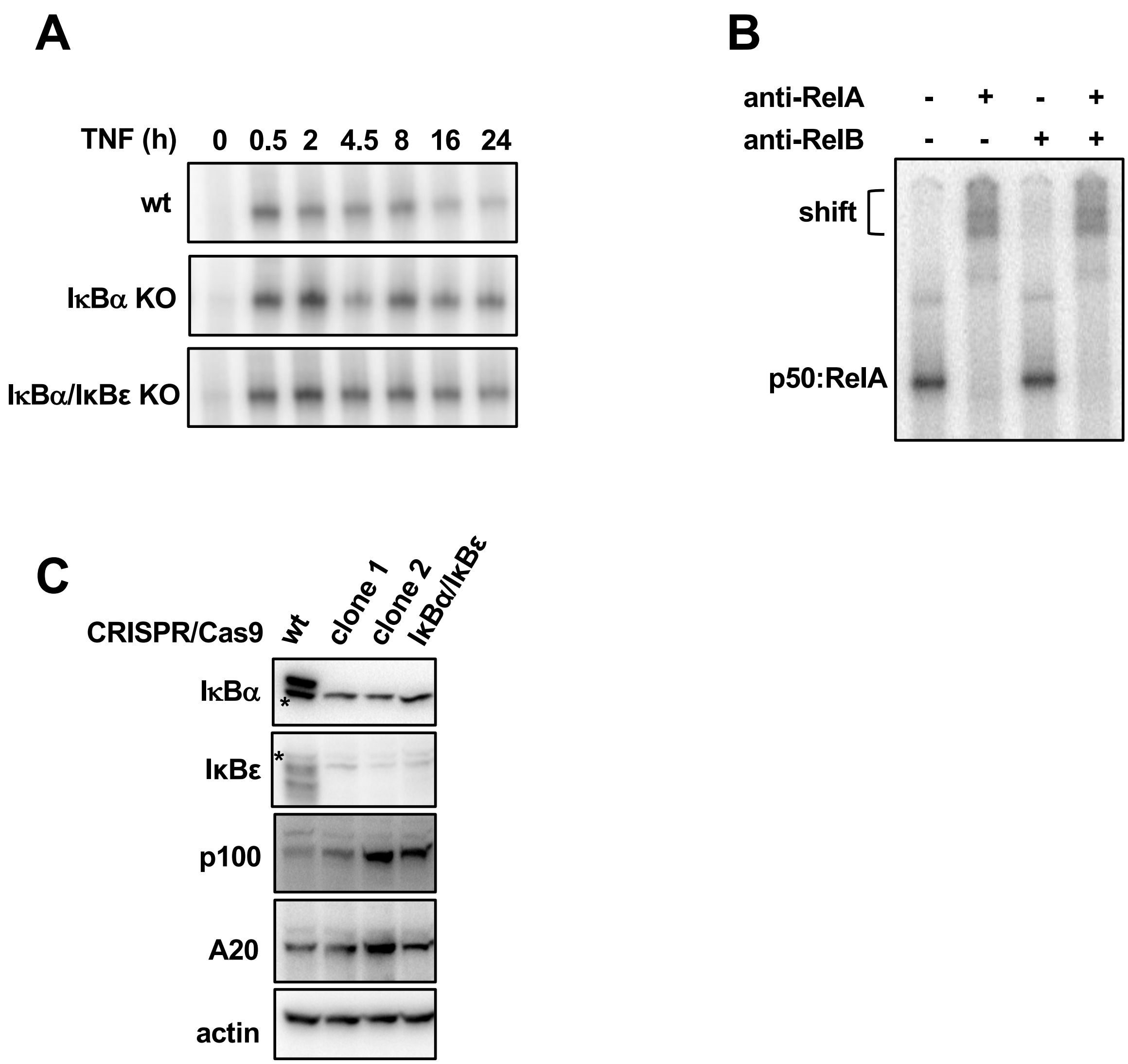

### Supplemental Figure 5

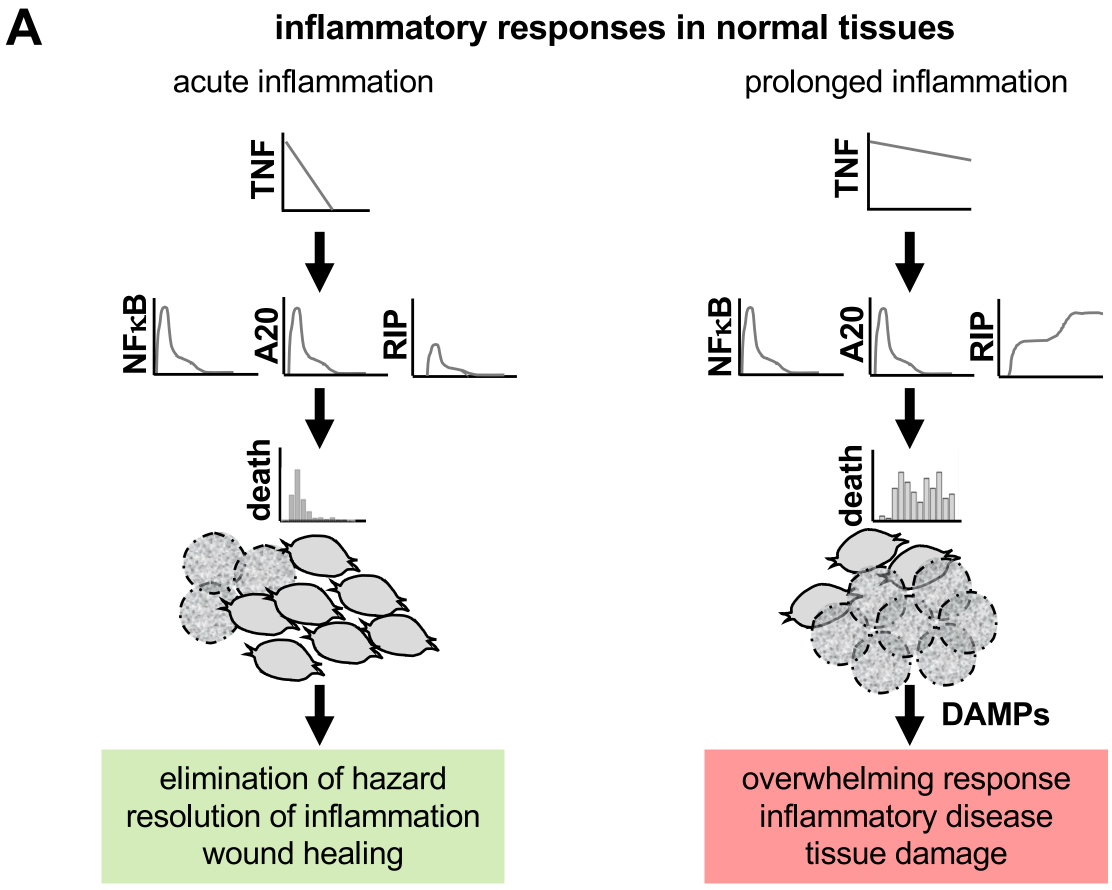


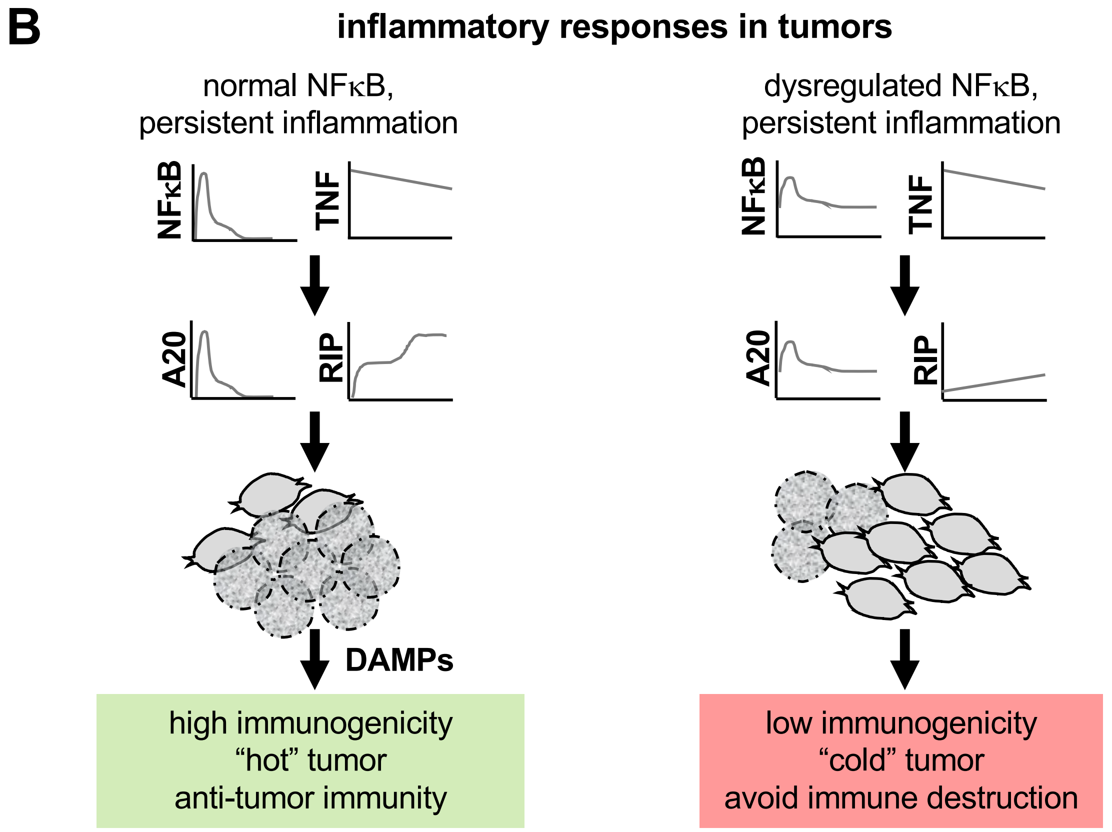

#### SUPPLEMENTARY FIGURE LEGENDS

##### Supplementary Figure 1

(A) Immune threats or tissue injury generate pathogen-associated molecular patterns (PAMPs) or damage-associated molecular patterns (DAMPs) to initiate inflammatory response. Tissue-resident macrophages respond with combined activation of NFκB and MAPK p38 pathways, and produce key inflammatory cytokines including IL6, IL1, and TNF. Stromal cells including fibroblasts perceive those cytokines and participate in response to inflammation. TNF-induced NFκB signaling drives release of chemokines (CCLs), matrix metalloproteinases (MMPs), and death ligands (FASL) to coordinate and resolve inflammatory cell infiltration. Alternatively, TNF may induce necroptotic cell death via RIP signaling, which generates more DAMPs to amplify inflammatory response. Whether TNF-induced NFκB cross-regulates necroptosis is unclear. (B) Proliferative index of L929 wildtype (wt) cells treated with TNF obtained from live-cell microscopy (mean of three independent experiments ± standard deviation). (C) Fractional cellular survival after 24 hours TNF in the presence or absence of fluorescent DNA staining (Hoechst and PI) quantified by crystal violet assay (mean of three independent experiments ± standard deviation; two-tailed Student's t-test revealed no statistically significant differences, n.s., $P>0.05$). (D) Fractional survival of TNF-treated L929 wt cells obtained via two independent methods (mean of three independent experiments ± standard deviation). (E) Immunoblot for pMLKL in L929 wt cells (representative data of three independent experiments). (F) Two-phased death rates in a clonal L929 wt cell population (mean of three independent experiments ± standard deviation). (G) TNF-induced NFκB activity in L929 wt cells measured via EMSA (representative data of three independent experiments). (H) Antibody against RelA, but not RelB induces shift of bands, revealing they largely consist of p50:RelA (representative data of three independent experiments). (I) Normalized RelA activity measured via EMSA and death rates in time course of TNF treatment (means of three independent experiments ± standard deviation). (J) Immunoblot for RelA in L929 wt and CRISP/Cas9 RelA-knockout (RelA KO) cell lines including clonal RelA KO population (asterisk indicates unspecific band present in all lanes). (K) Fractional survival after 24 hours of TNF treatment (mean of three independent experiments ± standard deviation; two-tailed Student's t-test **$P<0.01$). (L) Immunoblot for pMLKL (dash indicates empty lane; representative data of three independent experiments). (M) Representative images illustrating morphological characteristics of apoptosis vs. necrosis in RelA KO cells. Apoptotic cell (t1, left) shows cytoplasmic blebbing (t2) and delayed PI positivity of nucleus (t3) indicative of secondary necrosis. Necrotic cell (right, t1) shows cytoplasmic swelling and rapid PI positivity (t2). (N) Quantification of apoptotic vs. necrotic death in RelA KO cells manually curated from live-cell microscopy experiments using morphological criteria shown in (M) (200 cells analyzed per experiment; mean of three independent experiments ± standard deviation).

**Supplementary Figure 2**

Fractional survival quantified by crystal violet assay after treatment with TNF and (A) ROS inhibitor (ROSi), (B) JNK inhibitor (JNKi), and/or ZVAD (mean of three independent experiments ± standard deviation; two-tailed Student's t-test *P<0.05, **P<0.01, ***P<0.001, or no statistically significant difference, n.s., P>0.05). (C) TNF-induced mRNA expression measured via qRT-PCR (mean of three independent experiments ± standard deviation; two-tailed Student's t-test revealed no statistically significant results, P>0.05). (D) Immunoblot for FLIP-L (full) and its cleavage product (p43; representative data of three independent experiments). (E) Relative mRNA expression of A20 or cIAP2 measured via qRT-PCR, and normalized RelA activity measured via EMSA in wt cells treated with TNF (mean of three independent experiments ± standard deviation). (F) Immunoblot for A20 protein (representative data of three independent experiments). (G) Whole cell lysates depleted of RIPK3-containing protein complexes using anti-RIPK3 served as input for RIPK1 co-immunoprecipitation (IP) and subsequent immunoblot analysis. Flow through, FT. (H) Histogram of volume normalized mRNA copy numbers measured by smFISH for the NF$\kappa$B-responsive target genes A20 and I$\kappa$B$\alpha$ in wt cells treated with TNF (two independent experiments; third experiment shown in Figure 2D). (I) Immunoblot for cIAP2 using commercially available Pan-cIAP antibody and cIAP1-knockout cell line as control. As cIAP2 protein is below detection limit, CRISP/Cas9-mediated knockout was validated (J) via high resolution melt (HRM) analysis. (K) TNF-induced death rates obtained via live-cell microscopy (mean of three independent experiments ± standard deviation).

**Supplementary Figure 3**

(A) Detailed schematic diagram of model reaction network (for reactions and parameters see Tables in Supplementary Notes). (B-D) Model simulations (smoothed line represents population average, and shaded area the 20$^{th}$ percentile around the median) and experimental measurements (data points, mean of three independent experiments ± standard deviation) of indicated species over the time course of TNF treatment in L929 wildtype cells. A.U., arbitrary units. (E) Immunoblot for pMLKL and RIPK3 in detergent-soluble and -insoluble fractions of wildtype cells as a biochemical correlate of the active necrosome (representative data of three independent experiments). (F, G) Simulations and relative quantification of experimental measurements of necrosome activity in (E) (mean of three independent experiments ± standard deviation). (H) Immunoblot of A20 in wildtype (wt), parental RelA-knockout (KO) cells (-) or RelA KO cells expressing A20 from a constitutive transgene (pBabe-A20, +). Relative quantification across three independent experiments (mean ± standard deviation; two-tailed Student's t-test *$P<0.05$, or no statistically significant difference, n.s., $P>0.05$) (I) Probability of unimodal distributions of death times calculated by Hartigan's dip significance (mean of three independent experiments ± standard deviation; two-sample t-test revealed no statistical significance, n.s., $P>0.05$). (J) A20 immunoblot in wt or A20 KO cells (left). Simulated and measured distribution of death times in A20 KO cells (representative data of three independent experiments). (K) Immunoblot of A20 in wt and A20 KO cells reconstituted with an NFκB-inducible transgene (fIL8-A20), and relative quantification (mean of three independent experiments ± standard deviation). (L) Probability of unimodal distributions of death times calculated by Hartigan's dip significance in parental A20 KO cells (-) or A20 KO cells expressing fIL8-A20 (+). Mean of three independent experiments ± standard deviation; two-sample t-test **$P<0.01$). (M) 24-hour fractional survival in response to varying concentrations of TNF (mean of three independent experiments ± standard deviation).

**Supplementary Figure 4**

(A) TNF-induced NFκB activity in indicated cell lines measured via EMSA (representative data of three independent experiments). (B) Antibody for RelA, but not RelB induces shift of bands, revealing they largely consist of p50:RelA (representative data of three independent experiments). (C) Immunoblot for indicated proteins; clonal population 1 (clone 1) was selected for maintaining basal expression of p100 and A20 despite of IκBα/IκBε-knockout. Asterisks depict unspecific bands.

**Supplementary Figure 5**

(A) In normal tissues, the NFκB-A20-RIPK3 circuit protects the majority of fibroblasts from death induced by transient signals of TNF to participate in the coordination and resolution of acute inflammation (left). In conditions of prolonged inflammation, sustained TNF signals overcome the protective NFκB-A20-RIPK3 circuit, leading to massive necroptosis, the release of DAMPs, and overwhelming inflammation, which may contribute to inflammatory diseases and tissue damage (right). (B) In tumors with normal NFκB activity, sustained TNF signals may induce sufficient levels of necroptotic cell death to establish immunogenicity and an effective anti-tumor response (left). In contrast, in tumors with dysregulated NFκB activity, prolonged expression of A20 may contribute to low immunogenicity and avoidance of immune destruction (right).
