## Supplementary Notes for "An incoherent feedforward loop interprets NFκB/RelA dynamics to determine TNF-induced necroptosis decisions"

### Cell-death conceptual model

#### Model construction

To explore whether differential expression dynamics of a constitutive or stimulus-induced survival factor would produce differential time courses in the incidences of cell death, we constructed two conceptual models representing core network motifs of TNF-induced necroptosis as depicted in Figure 1. In the first, TNF induced activation of RIPK1/3 and the necroptosis effector pMLKL is counteracted by an unknown, constitutively expressed survival factor X. In the second model, TNF also induces IκB-controlled NFκB, which in turn induces the expression of the survival factor X. The modeling species and their initial values are listed in Table S1.

IκB-controlled NFκB activity is a coarse-grained version of previously published models (e.g. Werner et al. 2008), which we extended by adding inducible expression of survival factor X. The dynamics of each species were modeled by ordinary differential equations with synthesis and degradation terms (Kærn et al. 2003, Bintu et al. 2005, Krishna et al. 2006, Alon 2007, Ma et al., 2009, Tyson et al. 2010, Tang et al. 2017). The synthesis term contains the activation and inhibition as a product of Hill functions (MacArthur et al., 2009). The Hill functions have a sigmoidal shape suitable to represent the cascade-like activation of the protein kinases RIPK1/3 and pMLKL, which behave like a highly cooperative enzymatic reaction (Huang et al. 1996, Kapuy et al. 2009, Shu et al. 2013). The equations are stated below:

$$
\begin{aligned}
TNF &= \exp(-t/600),\\
\frac{d[RIP1]}{dt} &= k_{s1}\frac{(TNF/Km_1)^{n_1}}{(TNF/Km_1)^{n_1}+1}\frac{1}{([X]/Km_5)^{n_5}+1} - k_{d1}[RIP1],\\
\frac{d[pMLKL]}{dt} &= k_{s2}\frac{([RIP1]/Km_2)^{n_2}}{([RIP1]/Km_2)^{n_2}+1} - k_{d2}[pMLKL],\\
\frac{d[X]}{dt} &= -k_{d3}[X]/3,\\
\frac{d[IkB]}{dt} &= 0,\\
\frac{d[NFkB]}{dt} &= 0.
\end{aligned}
$$

$$
\begin{aligned}
TNF &= \exp(-t/600),\\
\frac{d[RIP1]}{dt} &= k_{s1}\frac{(TNF/Km_1)^{n_1}}{(TNF/Km_1)^{n_1}+1}\frac{1}{([X]/Km_5)^{n_5}+1} - k_{d1}[RIP1],\\
\frac{d[pMLKL]}{dt} &= k_{s2}\frac{([RIP1]/Km_2)^{n_2}}{([RIP1]/Km_2)^{n_2}+1} - k_{d2}[pMLKL],\\
\frac{d[X]}{dt} &= k_{s3}\frac{([NFkB]/Km_3)^{n_3}}{([NFkB]/Km_3)^{n_3}+1} - k_{d3}[X],\\
\frac{d[IkB]}{dt} &= k_{s4}\frac{([NFkB]/Km_4)^{n_4}}{([NFkB]/Km_4)^{n_4}+1}\frac{1}{(TNF/Km_6)^{n_6}+1} - k_{d4}[IkB],\\
\frac{d[NFkB]}{dt} &= k_{s5}\frac{1}{([IkB]/Km_7)^{n_7}+1} - k_{d5}[NFkB].
\end{aligned}
$$

The parameters include synthesis rates $k_s$, degradation rates $k_d$, Hill coefficients $n$, and Km values in the Hill functions. The synthesis and degradation rates determine the activation and decay rate of the curves of protein concentration; Hill coefficients correspond to the sensitivity upon activation; and Km values denote the dependence of activation on the concentration of activator or inhibitor. We assumed that all the parameters are constant, as listed in Table S2. The unit of time in the simulation is minute. To account for cell-to-cell heterogeneity in expression of survival factor X (Friedman et al. 2006, Shalek et al. 2013), we distributed either its constitutive expression in the first model, or the NFκB-responsive synthesis rate in the second model. The distributed parameters are listed in Table S3.

#### Model simulations

##### Death time distribution

For each model, we simulated 300 single cells, with each simulation sampling one set of values from the parameter distributions. The irreversible cell death event corresponds to the death marker pMLKL reaching a threshold (Spencer et al. 2009), and the death time is registered as the time of threshold crossing. The conceptual modeling framework can be dimensionless, and we added units that are consistent with the more extensive model of necroptosis below. Since only the relative value of protein to the threshold determines the cell death time, the protein values were normalized within a certain range with arbitrary units (A.U.). We conducted these simulations using MATLAB® file DeathConcept2.m. All simulations were done using MATLAB® version R2019a.

In Figure 1, we plotted the time course of cells undergoing death by hourly binning the number of simulations in which pMLKL exceeds the threshold. While the first model returns a unimodal, the second model returns a non-unimodal distribution of death times.

##### Parameter scan

These two conceptual models aim to provide a coarse-grained description of necroptosis kinetics. As we aimed to compare relative differences in necroptosis kinetics between both versions depending on whether constitutive or stimulus-induced survival factors are present, the absolute parameter values were not critical. However, the dynamics of IκB-controlled NFκB were adjusted to coarsely match previously published models (Werner et al. 2008, Krishna et al. 2006). To implement dependence of pMLKL on the stimulus and protective protein, we further adjusted the dynamics of upstream proteins. For example, we chose Km values in the range of the concentration of the activators or inhibitors during their transient dynamics, such that the activation or inhibition took effect when the protein accumulates to the Km values. The synthesis and degradation rates of upstream proteins were set accordingly to ensure the proteins reached Km values to affect the downstream proteins during the time window of simulation. We allowed the effective Hill coefficients in the range of [1,5] (Goldbeter et al. 1981, Huang et al. 1996). The Hill coefficient was adjusted to tune the sensitivity of the activation or inhibition: if downstream protein was sharply affected by the activator or inhibitor, the relatively large Hill coefficient was implied.

In order to test the dependence of cell death time distributions on the parameters, we performed parameter sensitivity analysis for the synthesis rates, degradation rates and Km values of all reactions. For each simulation, we sampled one set of parameters from the parameter regime of 2-fold higher and 2-fold lower relative to the original values above, as in Table S4. We then performed statistical analysis of the distributions of death times to calculate a probability of unimodal distributions as explained in the Methods section (Hartigan et al. 1985). We classified the cell death time distribution as unimodal or non-unimodal by separating the unimodality score at the threshold 0.5.

We repeated the simulation 1000 times for each model, and classified the returned death time distributions into 3 types: (1) “survival ratio > 40%” with overall survival of > 40% of the cell population, (2) “unimodal” with overall survival of < 40% and a unimodality score of > 0.5, (3) “non-unimodal” with overall survival of < 40% and a unimodality score of < 0.5. We excluded simulations with > 40% survival to increase the confidence in classifying the modality of death time distribution. The simulations showed that the first model majorly returns unimodal distributions, whereas the second model preferably generates non-unimodal distributions of death times:

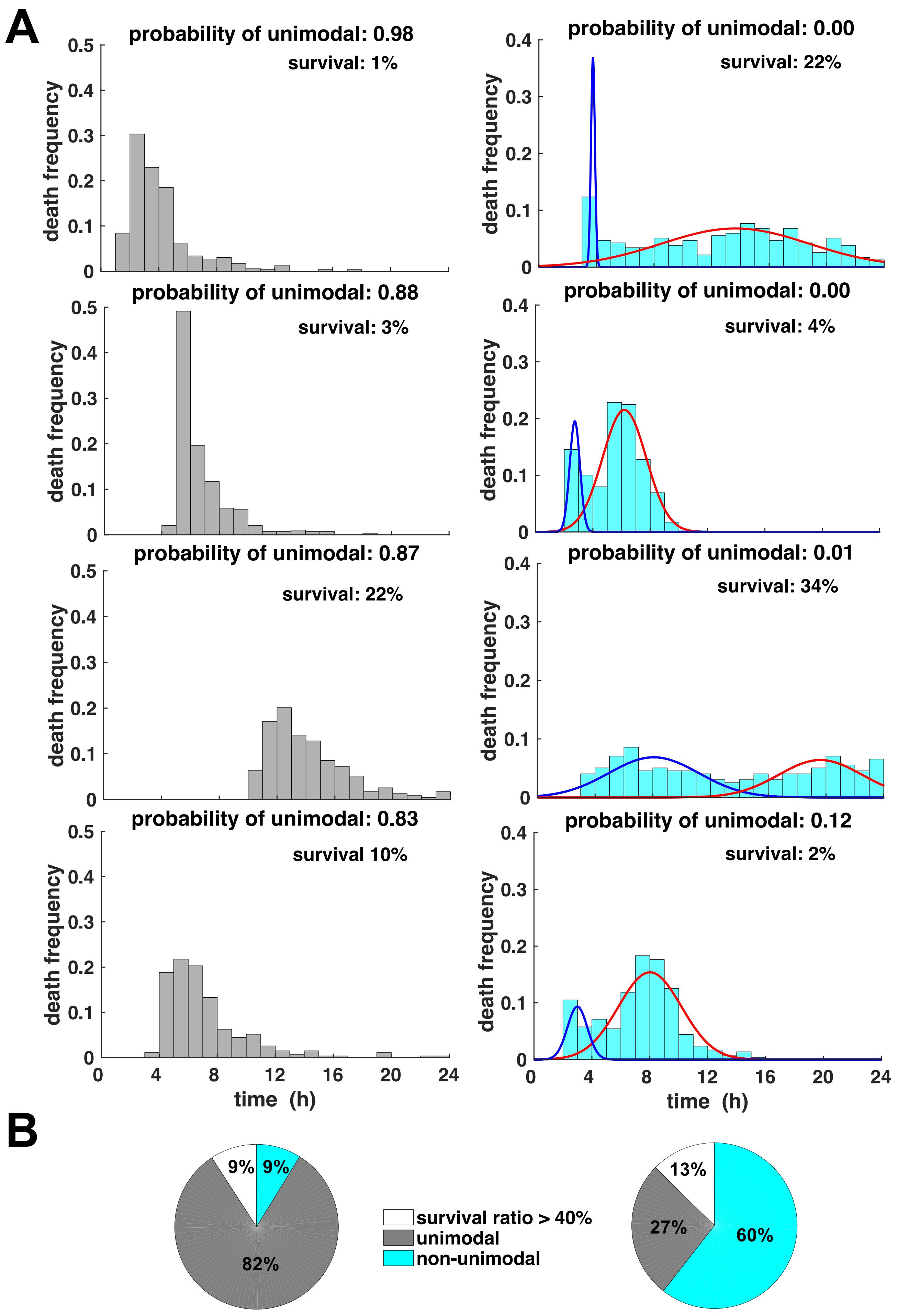

(A) Representative simulations of death times after parameter sensitivity analysis for both conceptual modeling versions of TNF-induced necroptosis (left: constitutive, right: inducible survival factor). Probability of unimodality calculated by Hartigan's dip significance test, and classified as unimodal or non-unimodal at threshold 0.5. (B) Percentages of simulations revealing unimodal or non-unimodal distributions of death times; simulations with >40% overall survival were excluded.

### Mathematic model for TNF-induced necroptosis

#### Model construction

To construct a mechanistic model of NF$\kappa$B and necroptosis network dynamics, we first combined the previously published model of TNFR signaling to NF$\kappa$B (Werner et al. 2008) with the 4-IκB-containing NF$\kappa$B module (Shih et al. 2009). This merged version reflects TNF-induced activation of complex I to activate IKK, which causes the degradation of IκB proteins and activation of NF$\kappa$B to initiate transcription of IκBs as well as A20, with A20 counteracting the activation of complex I (Supplementary Figure 3A). In a next step, we extended this model by adding a course-grained necroptosis module depicting complex I activating RIPK1, RIPK3 and the effector pMLKL, with A20 inhibiting the activation of RIPK3 (Supplementary Figure 3A). While this model of necroptosis is limited to the core biochemical signaling processes, it was sufficient to recapitulate TNF-induced signaling kinetics controlled by NF$\kappa$B activity dynamics.

##### TNFR-NF$\kappa$B-4-IκB module

We adapted the module of TNF signaling to NF$\kappa$B via IKK from [Werner et al. 2008 G&D], which includes the negative feedback regulators IκB$\alpha$, IκBε, and IκBβ as well as the regulator A20. For completeness, we have listed all species, reactions, initial values and the parameters in Table S5 and Table S6. We then added the 4-IκB-containing NF$\kappa$B module from (Shih et al. 2009) to include IκB$\delta$ inhibitory activity. We listed all species, reactions, initial values and parameters in Table S5 and Table S6 as well. To parameterize this model to L929 cells, we used experimental measurements of NFκB activity (EMSA) and IκB expression (Immunoblot). TNF half-life was set to 10 hours to match our data. All changes are listed in Table S7.

##### Necroptosis module

To quantitatively understand the kinetics of necroptosis signaling, we constructed a minimal model to describe the dynamics of the three molecular species RIPK1, RIPK3, and pMLKL (active forms of all regulators). Each equation is composed of synthesis and decay terms (Kærn et al. 2003, Bintu et al. 2005, Alon 2007, Shu et al. 2013, Ma et al., 2009, Tyson et al. 2010, Tang et al. 2017). Similar as the model in Fig.1, the synthesis term is a product of activation and inhibition by other signaling proteins following the schematic, and modeled by a product of Hill functions (MacArthur et al. 2009). The inhibition of necroptosis by A20 is also modeled by a Hill function in the synthesis term from RIP1 to RIP3. The equations are stated below:

$$\frac{d[RIP1]}{dt} = k_{s1}\frac{[([C1]+[C1_off])/Km_1]^{n_1}}{[([C1]+[C1_off])/Km_1]^{n_1}+1} - k_{d1}[RIP1],$$
$$\frac{d[RIP3]}{dt} = k_{s2}\frac{([RIP1]/Km_2)^{n_2}}{([RIP1]/Km_3)^{n_2}+1}\frac{1}{([A20]/Km_3)^{n_3}+1} - k_{d2}[RIP3],$$
$$\frac{d[pMLKL]}{dt} = k_{s3}\frac{([RIP3]/Km_4)^{n_4}}{([RIP3]/Km_4)^{n_4}+1} - k_{d3}[pMLKL].$$

The parameters contain synthesis rates, degradation rates, Hill coefficients, and Km values. They were fit to experimental measurements of biochemical markers of necroptosis (pMLKL and RIPK3 measurements via Western blot), death kinetics (microscopy) and fractional survival (microscopy and/or crystal violet assay). For example, the synthesis and degradation rates were determined by the activation rate of the time course data of RIPK1, RIPK3 and pMLKL; Hill coefficients were adjusted to fit the steepness of the activation curve upon stimulation; and Km values were chosen to match the dependence on the concentration of activator or inhibitor. We repeated this fitting procedure until the simulated time course of signaling proteins is fitted with their measured activity at different time points, with the parameters in Table S8.

#### Model simulations

##### Wildtype

We simulated 300 cells comparable to each microscopy experiment, and simulated the time course of each molecular species in the model. To account for cell-to-cell heterogeneity, we distributed a minimal number of parameters: RIPK1 to RIPK3 activation rate, which obeys a unimodal distribution (equivalent to unknown constitutive survival mechanisms); and A20 mRNA synthesis rate, which follows a bimodal distribution in line with our single-molecule FISH measurements. All the distributed parameters are listed in Table S9. We used the population average of simulated mRNA and/or protein concentration to fit with experimental measurements for NFκB, IκB, A20, RIPK3 and pMLKL. We plotted the simulated time course along with experimental measurements for key molecules, such as nuclear NFκB, A20 mRNA, A20 protein, RIP3, and pMLKL (Supplementary Figure 3B-G). The shaded area is the range of the 20th percentile around the median value of the simulated 300 cells. For each species, we first normalized the data to simulation, since the simulation has the protein in the unit of μM and time in the unit of minute. After fitting to the data, we renormalized both the data and simulation back to value of experiment, such that the figures are in the same scale of original normalized data under an arbitrary unit (A.U.).

For each simulated cell, we tracked its death event by using a threshold for pMLKL, which we also adopted for the conceptual model. We then plotted the death time distributions, death rates and fractional survival in Figure 3B, C. Using the distributed parameters, the repeated simulations of single cells recapitulated the bimodal death time distributions, two-phased death rates and fractional survival of experimental measurements. The simulations were executed in the MATLAB® script NecroptosisMain1.m.

##### RelA-knockout

After validating that the model accurately recapitulated the time course of key signaling molecules, as well as the death time distribution of wildtype cells, we applied it to simulate the scenario of NFκB/RelA-knockout cells (RelA KO). We modeled RelA KO by setting the value of NFκB (RelA) to zero. Without RelA, TNF did not induce A20 expression, and therefore cell death occurred more rapidly within the early phase of the simulated time course. Accordingly, the simulation returned unimodal death time distributions, single-phased death rates and matched overall fractional survival of experimental measurements. The simulated time course is plotted in Figure 3B, C. We concluded that the model can accurately recapitulate the RelA-dependent regulation of necroptosis kinetics.

##### A20-knockout

Similarly, we modeled the scenario of A20-knockout (A20 KO) by setting A20 mRNA to zero. Without the protection of A20 protein, cell death occurred rapidly within the early phase of the simulated time course, and followed a unimodal shape, which was in line with measurements (Supplementary Figure 3I). It demonstrated that our model properly fits the strength of A20 inhibition on cell death.

##### Constitutive A20 expression

Next, we explored whether constitutive or transient A20 expression is crucial for the regulation of necroptosis kinetics. After setting TNF-inducible expression of A20 to zero, we simulated 24 hours of TNF treatment with constitutive A20 transcription rates modified by factors of $2^x$ ($x$ = -1, 0, 1; “basal / RelA-knockout” resembled by $x$ = -1). We simulated 300 cells per condition and plotted the resulting death time distributions as death count over time (Figure 3D).

##### TNF pulse stimulation of different durations

We applied the model to simulate overall fractional survival after different duration doses of TNF (10 ng/ml). By increasing the decay rate of TNF (1500-fold) at 3, 6, 12, or 24 hours, TNF soon reaches zero after those time points, respectively. As TNF duration increased, the simulated fractional survival of wildtype or RelA-knockout cells at 24 hours decreased (Figure 3I). The model predictions were tested experimentally by washing out TNF at chosen time points (Figure 3J).

##### Synthetically regulated NFκB

In order to explore how the duration of A20 expression controls overall fractional survival in response to TNF, we placed the A20 promoter under the control of a “synthetic” NFκB activity. Synthetically regulated NFκB activity was implemented using a pulse function, which is not restrained by IκB feedback regulators and serves as an input to control mA20 expression. The pulse function of NFκB activity started at time zero with 3 nM, and increased to be 83 nM, and we tested a set of durations: 0.5, 1, 2, 4, 8, 16 hours. Correspondingly, the simulated induction of mA20 followed similar dynamics, as shown in Figure 4A, B. As the duration of synthetic NFκB activity and subsequent mA20 expression increased, the fractional survival at 24 hours increased as well. We validated this prediction experimentally by using IκBα/IκBε-knockout cells showing elevated NFκB activity and prolonged expression of mA20 (Figure 4C-G). These findings suggested that dysregulated activation of NFκB and inducible A20 may protect cells from cytotoxic functions of long-term TNF-exposure.

### Supplementary Tables for cell-death conceptual models

#### Supplementary Table S1: Model species for cell death conceptual model

| | Model Species | Initial value in Model 1 | Initial value in Model 2 |
|---|---|---|---|
| **1** | RIP1/3 | 0 | 0 |
| **2** | pMLKL | 0 | 0 |
| **3** | X | 0.05*lognormal(0,0.3) μM, where lognormal(a,b) is a log-normal distribution with mean a and variable b. | 0 |
| **4** | IκB | 0 | 0 |
| **5** | NF-κB | 0 | 0.01 μM |

#### Supplementary Table S2: Parameters for cell death conceptual models

| # | Reaction | Parameter | Parameter Value | Model 1 | Model 2 |
|---|---|---|---|---|---|
| | TNF decay | Half life | 600 min | Yes | Yes |
| 1 | TNF --> RIP1/3, X --\| RIP1/3 | Synthesis rate $k_{s1}$ | 0.1 min$^{-1}$ | Yes | Yes |
| 2 | RIP1/3 --> pMLKL | Synthesis rate $k_{s2}$ | 0.1 min$^{-1}$ | Yes | Yes |
| 3 | NFkB –> X | Synthesis rate $k_{s3}$ | 5 min$^{-1}$ | No | Yes |
| 4 | TNF --\| IkB, NFkB --> IkB | Synthesis rate $k_{s4}$ | 5 min$^{-1}$ | No | Yes |
| 5 | IkB --\| NFkB | Synthesis rate $k_{s5}$ | 1 min$^{-1}$ | No | Yes |
| 1 | RIP1/3 => | Degradation rate $k_{d1}$ | 0.01 μM$^{-1}$ min$^{-1}$ | Yes | Yes |
| 2 | pMLKL => | Degradation rate $k_{d2}$ | 0.1 μM$^{-1}$ min$^{-1}$ | Yes | Yes |
| 3 | X => | Degradation rate $k_{d3}$ | 1 μM$^{-}$ min$^{-1}$ | No | Yes |
| 4 | IkB => | Degradation rate $k_{d4}$ | 1 μM$^{-1}$ min$^{-1}$ | No | Yes |
| 5 | NFkB => | Degradation rate $k_{d5}$ | 1 μM$^{-1}$ min$^{-1}$ | No | Yes |
| 1 | TNF --> RIP1/3 | $Km_1$ | 1 μM | Yes | Yes |
| 2 | RIP1/3 --> pMLKL | $Km_2$ | 0.15 μM | Yes | Yes |
| 3 | NFkB –> X | $Km_3$ | 0.5 μM | No | Yes |
| 4 | NFkB –> IkB | $Km_4$ | 0.3 μM | No | Yes |
| 1 | X --\| RIP1/3 | $Km_5$ | 0.3 μM | Yes | Yes |
| 4 | TNF --\| IkB | $Km_6$ | 0.3 μM | No | Yes |
| 5 | IkB --\| NFkB | $Km_7$ | 0.03 μM | No | Yes |
| 1 | TNF --> RIP1/3 | Hill coefficient $n_1$ | 3 | Yes | Yes |
| 2 | RIP1/3 --> pMLKL | Hill coefficient $n_2$ | 3 | Yes | Yes |
| 3 | NFkB –> X | Hill coefficient $n_3$ | 3 | No | Yes |
| 4 | NFkB –> IkB | Hill coefficient $n_4$ | 3 | No | Yes |
| 1 | X --\| RIP1/3 | Hill coefficient $n_5$ | 3 | Yes | Yes |
| 4 | TNF --\| IkB | Hill coefficient $n_6$ | 3 | No | Yes |
| 5 | IkB --\| NFkB | Hill coefficient $n_7$ | 3 | No | Yes |

#### Supplementary Table S3: Distributed parameters for cell death conceptual model

| # | Parameter | Parameter Value | Parameter distribution |
|---|---|---|---|
| 3 | Synthesis rate $k_{s3}$ | Distributed | 0.5*normal(7,8)+ 0.5*normal(14,2), where normal(a,b) is a normal distribution with mean a and variable b. |

#### Supplementary Table S4: Perturbed range of parameters for cell death conceptual model

| # | Parameter | Parameter Value | Sampling range |
|---|---|---|---|
| 1 | Synthesis rate $k_{s1}$ | 0.1 min$^{-1}$ | [0.5, 1, 2] |
| 2 | Synthesis rate $k_{s2}$ | 0.1 min$^{-1}$ | [0.5, 1, 2] |
| 3 | Synthesis rate $k_{s3}$ | 5 min$^{-1}$ | [0.5, 1, 2] |
| 4 | Synthesis rate $k_{s4}$ | 5 min$^{-1}$ | [0.5, 1, 2] |
| 5 | Synthesis rate $k_{s5}$ | 1 min$^{-1}$ | [0.5, 1, 2] |
| 1 | Degradation rate $k_{d1}$ | 0.01 μM$^{-1}$ min$^{-1}$ | [0.5, 1, 2] |
| 2 | Degradation rate $k_{d2}$ | 0.1 μM$^{-1}$ min$^{-1}$ | [0.5, 1, 2] |
| 3 | Degradation rate $k_{d3}$ | 1 μM$^{-1}$ min$^{-1}$ | [0.5, 1, 2] |
| 4 | Degradation rate $k_{d4}$ | 1 μM$^{-1}$ min$^{-1}$ | [0.5, 1, 2] |
| 5 | Degradation rate $k_{d5}$ | 1 μM$^{-1}$ min$^{-1}$ | [0.5, 1, 2] |
| 1 | $Km_1$ | 1 μM | [0.5, 1, 2] |
| 2 | $Km_2$ | 0.15 μM | [0.5, 1, 2] |
| 3 | $Km_3$ | 0.5 μM | [0.5, 1, 2] |
| 4 | $Km_4$ | 0.3 μM | [0.5, 1, 2] |
| 1 | $Km_5$ | 0.3 μM | [0.5, 1, 2] |
| 4 | $Km_6$ | 0.3 μM | [0.5, 1, 2] |
| 5 | $Km_7$ | 0.03 μM | [0.5, 1, 2] |

### Supplementary Tables for TNF-induced necroptosis model

#### Supplementary Table S5: Model species

| | Model Species | Model Nomenclature | Initial μM | Location |
|---|---|---|---|---|
| **1** | IκBα | IkBa | 0 | Cytoplasm |
| **2** | IκBα | IkBan | 0 | Nucleus |
| **3** | IκBα-NFκB | IkBaNFkB | 0 | Cytoplasm |
| **4** | IκBα-NFκB | IkBaNFkBn | 0 | Nucleus |
| **5** | IκBα mRNA | IkBat | 0 | Cytoplasm |
| **6** | IκBβ | IkBb | 0 | Cytoplasm |
| **7** | IκBβ | IkBbn | 0 | Nucleus |
| **8** | IκBβ-NFκB | IkBbNFkB | 0 | Cytoplasm |
| **9** | IκBβ-NFκB | IkBbNFkBn | 0 | Nucleus |
| **10** | IκBϵ mRNA | IkBbt | 0 | Cytoplasm |
| **11** | IκBϵ | IkBe | 0 | Cytoplasm |
| **12** | IκBϵ | IkBen | 0 | Nucleus |
| **13** | IκBϵ-NFκB | IkBeNFkB | 0 | Cytoplasm |
| **14** | IκBϵ-NFκB | IkBeNFkBn | 0 | Nucleus |
| **15** | IκBϵ mRNA | IkBet | 0 | Cytoplasm |
| **16** | IκBδ | IkBd | 0 | Cytoplasm |
| **17** | IκBδ | IkBdn | 0 | Nucleus |
| **18** | IκBδ-NFκB | IkBdNFkB | 0 | Cytoplasm |
| **19** | IκBδ-NFκB | IkBdNFkBn | 0 | Nucleus |
| **20** | IκBδ mRNA | IkBdt | 0 | Cytoplasm |
| **21** | NF-κB | NFkB | 0 | Cytoplasm |
| **22** | NF-κB | NFkBn | 0.125 | Nucleus |
| **23** | TAK1 (inactive) | IKKK_off | 0.1 | Cytoplasm |
| **24** | TAK1 (active) | IKKK | 0 | Cytoplasm |
| **25** | IKK (inactive) | IKK_off | 0.1 | Cytoplasm |
| **26** | IKK (active) | IKK | 0 | Cytoplasm |
| **27** | IKK (auto-inactivated) | IKK_i | 0 | Cytoplasm |
| **28** | TNF | tnf | 0 | Extracellular |
| **29** | TNF Receptor Monomer | tnfrm | 0 | Cell Surface |
| **30** | TNF Receptor Trimer | TNFR | 0 | Cell Surface |
| **31** | TNF-Bound TNF Receptor Trimer | TNFRtnf | 0 | Cell Surface |
| **32** | TNFR Complex I (active) | C1 | 0 | Cell Surface |
| **33** | TNFR Complex I (inactive) | C1_off | 0 | Cell Surface |
| **34** | TNF-Bound TNFR Complex I (active) | C1tnf | 0 | Cell Surface |
| **35** | TNF-Bound TNFR Complex I (inactive) | C1_tnf_off | 0 | Cell Surface |
| **36** | TRAF-TRADD-RIP | TTR | 8.3e-4 | Cytoplasm |
| **37** | A20 | A20 | 0 | Cytoplasm |
| **38** | A20 mRNA | A20t | 0 | Cytoplasm |
| **39** | RIP3 | RIP3 | 0 | Cytoplasm |
| **40** | pMLKL | pMLKL | 0 | Cytoplasm |
| **41** | RIP1 | RIP1 | 0 | Cytoplasm |

Note for Supplementary Table S5: There are 41 species in the model. Each is represented by a unique name (nomenclature), with an initial concentration and cellular localization.

#### Supplementary Table S6: Model reactions

| NFκB Activation Module | | | | | |
|---|---|---|---|---|---|
| ***IκB mRNA and Protein Synthesis Reactions*** | | | | | |
| **#** | **Reaction** | **Parameter Value** | **Category** | **Location** | **Source of Parameter Value** |
| **1** | => IkBat (constitutive) | 7 E-5 $min^{-1}$ | RNA Synth. | - | Parameter value chosen to fit mRNA and protein expression profiles as measured by RNase Protection (RPA) and Western Blot assays. |
| **2** | => IkBbt (constitutive) | 1 E-5 $min^{-1}$ | RNA Synth. | - | Refer to #1. |
| **3** | => IkBet (constitutive) | 1 E-6 $min^{-1}$ | RNA Synth. | - | Refer to #1. |
| **72** | => IkBdt (constitutive) | 1 E-7 min-1 | RNA Synth. | - | Refer to #1. |
| **4**<br>**7**<br>**10** | => IkBat (induced by NFkBn) | 8 $\mu M^{-2}$ $min^{-1}$<br>Hill Coefficient: 3.0<br>Delay: 0 min | RNA Synth. | - | (Werner et al., 2005)<br>(Werner et al., 2005)<br>(Kearns et al., 2006) and unpublished results |
| **5**<br>**8**<br>**11** | => IkBbt (induced by NFkBn) | 0.02 $\mu M^{-2}$ $min^{-1}$<br>Hill Coefficient: 3.0<br>Delay: 37 min | RNA Synth. | - | (Kearns et al., 2006)<br>(Werner et al., 2005)<br>(Kearns et al., 2006) and unpublished results |
| **6**<br>**9**<br>**12** | => IkBet (induced by NFkBn) | 0.3 $\mu M^{-2}$ $min^{-1}$<br>Hill Coefficient: 3.0<br>Delay: 37 min | RNA Synth. | - | (Kearns et al., 2006)<br>(Werner et al., 2005)<br>(Kearns et al., 2006) and unpublished results |
| **73**<br>**74**<br>**75** | => IkBdt (induced by NFkBn) | 0.025 $\mu M^{-2}$ $min^{-1}$<br>Hill Coefficient: 3.0<br>Delay: 90 min | RNA Synth. | - | (Shih et al., 2009) |
| **13** | IkBat => | 0.035 $min^{-1}$ | RNA Deg. | Cytoplasm | mRNA half-life measurements using actinomycin-D treatment of cells and RPA. (unpublished results) |
| **14** | IkBbt => | 3 E-3 $min^{-1}$ | RNA Deg. | Cytoplasm | Refer to #7. |
| **15** | IkBet => | 4 E-3 $min^{-1}$ | RNA Deg. | Cytoplasm | Refer to #7. |
| **76** | IkBdt => | 2 E-3 $min^{-1}$ | RNA Deg. | Cytoplasm | Refer to #7. |
| **16** | => IkBa | 0.25 $min^{-1}$ | Prot. Synth. | Cytoplasm | (Hoffmann et al., 2002) |

| | | | | | |
|---|---|---|---|---|---|
| **17** | => IkBb | 0.25 $min^{-1}$ | Prot. Synth. | Cytoplasm | (Hoffmann et al., 2002) |
| **18** | => IkBe | 0.25 $min^{-1}$ | Prot. Synth. | Cytoplasm | (Hoffmann et al., 2002) |
| **77** | => IkBd | 0.25 $min^{-1}$ | Prot. Synth. | Cytoplasm | (Shih et al., 2009) |
| ***IκB and NFκB Cellular Localization Reactions*** | | | | | |
| **19** | IkBa => IkBan | 0.09 $min^{-1}$ | Import | - | (Werner et al., 2005) |
| **20** | IkBb => IkBbn | 0.009 $min^{-1}$ | Import | - | (Werner et al., 2005) |
| **21** | IkBe => IkBen | 0.045 $min^{-1}$ | Import | - | (Werner et al., 2005) |
| **78** | IkBd => IkBdn | 0.045 $min^{-1}$ | Import | - | (Shih et al., 2009) |
| **22** | NFkB => NFkBn | 5.4 $min^{-1}$ | Import | - | (Werner et al., 2005) |
| **23** | IkBan => IkBa | 0.012 $min^{-1}$ | Export | - | (Werner et al., 2005) |
| **24** | IkBbn => IkBb | 0.012 $min^{-1}$ | Export | - | (Werner et al., 2005) |
| **25** | IkBen => IkBe | 0.012 $min^{-1}$ | Export | - | (Werner et al., 2005) |
| **79** | IkBdn => IkBd | 0.012 $min^{-1}$ | Export | - | (Shih et al., 2009) |
| **26** | NFkBn => NFkB | 0.0048 $min^{-1}$ | Export | - | (Werner et al., 2005) |
| **27** | IkBaNFkB => IkBaNFkBn | 0.276 $min^{-1}$ | Import | - | (Werner et al., 2005) |
| **28** | IkBbNFkB => IkBbNFkBn | 0.0276 $min^{-1}$ | Import | - | (Werner et al., 2005) |
| **29** | IkBeNFkB => IkBeNFkBn | 0.138 $min^{-1}$ | Import | - | (Werner et al., 2005) |
| **80** | IkBdNFkB => IkBdNFkBn | 0.276 $min^{-1}$ | Import | - | (Shih et al., 2009) |
| **30** | IkBaNFkBn => IkBaNFkB | 0.828 $min^{-1}$ | Export | - | (Werner et al., 2005) |
| **31** | IkBbNFkBn => IkBbNFkB | 0.414 $min^{-1}$ | Export | - | (Werner et al., 2005) |
| **32** | IkBeNFkBn => IkBeNFkB | 0.414 $min^{-1}$ | Export | - | (Werner et al., 2005) |
| **81** | IkBdNFkBn => IkBdNFkB | 0.414 $min^{-1}$ | Export | - | (Shih et al., 2009) |
| ***IκB Protein Degradation Reactions*** | | | | | |
| **33** | IkBa => | 0.12 $min^{-1}$ | Prot. Deg. | Cytoplasm | (O'Dea et al., 2007) |
| **34** | IkBb => | 0.18 $min^{-1}$ | Prot. Deg. | Cytoplasm | (O'Dea et al., 2007) |
| **35** | IkBe => | 0.18 $min^{-1}$ | Prot. Deg. | Cytoplasm | (O'Dea et al., 2007) |
| **82** | IkBd => | 1.4E-3 $min^{-1}$ | Prot. Deg. | Cytoplasm | (Shih et al., 2009) |
| **36** | IkBan => | 0.12 $min^{-1}$ | Prot. Deg. | Nucleus | (O'Dea et al., 2007) |
| **37** | IkBbn => | 0.18 $min^{-1}$ | Prot. Deg. | Nucleus | (O'Dea et al., 2007) |
| **38** | IkBen => | 0.18 $min^{-1}$ | Prot. Deg. | Nucleus | (O'Dea et al., 2007) |
| **93** | IkBdn => | 1.4E-3 $min^{-1}$ | Prot. Deg. | Nucleus | (Shih et al., 2009) |
| **39** | IkBaNFkB => NFkB | 6E-5 $min^{-1}$ | Prot. Deg. | Cytoplasm | (O'Dea et al., 2007) |
| **40** | IkBbNFkB => NFkB | 6E-5 $min^{-1}$ | Prot. Deg. | Cytoplasm | (O'Dea et al., 2007) |
| **41** | IkBeNFkB => NFkB | 6E-5 $min^{-1}$ | Prot. Deg. | Cytoplasm | (O'Dea et al., 2007) |
| **84** | IkBdNFkB => NFkB | 6E-5 $min^{-1}$ | Prot. Deg. | Cytoplasm | (Shih et al., 2009) |
| **42** | IkBaNFkBn => NFkBn | 6E-5 $min^{-1}$ | Prot. Deg. | Nucleus | (O'Dea et al., 2007) |
| **43** | IkBbNFkBn => NFkBn | 6E-5 $min^{-1}$ | Prot. Deg. | Nucleus | (O'Dea et al., 2007) |
| **44** | IkBeNFkBn => NFkBn | 6E-5 $min^{-1}$ | Prot. Deg. | Nucleus | (O'Dea et al., 2007) |
| **85** | IkBdNFkBn => NFkBn | 6E-5 $min^{-1}$ | Prot. Deg. | Nucleus | (Shih et al., 2009) |
| ***IκB:NFκB Association and Dissociation Reactions*** | | | | | |
| **45** | IkBa + NFkB => IkBaNFkB | 30 $\mu M^{-1}$ $min^{-1}$ | Association | Cytoplasm | (Hoffmann et al., 2002) |
| **46** | IkBb + NFkB => IkBbNFkB | 30 $\mu M^{-1}$ $min^{-1}$ | Association | Cytoplasm | (Hoffmann et al., 2002) |
| **47** | IkBe + NFkB => IkBeNFkB | 30 $\mu M^{-1}$ $min^{-1}$ | Association | Cytoplasm | (Hoffmann et al., 2002) |
| **86** | IkBd + NFkB => IkBdNFkB | 30 $\mu M^{-1}$ $min^{-1}$ | Association | Cytoplasm | (Shih et al., 2009) |
| **48** | IkBan + NFkBn => IkBaNFkBn | 30 $\mu M^{-1}$ $min^{-1}$ | Association | Nucleus | (Hoffmann et al., 2002) |
| **49** | IkBbn + NFkBn => IkBbNFkBn | 30 $\mu M^{-1}$ $min^{-1}$ | Association | Nucleus | (Hoffmann et al., 2002) |
| **50** | IkBen + NFkBn => IkBeNFkBn | 30 $\mu M^{-1}$ $min^{-1}$ | Association | Nucleus | (Hoffmann et al., 2002) |
| **87** | IkBdn + NFkBn => IkBdNFkBn | 30 $\mu M^{-1}$ $min^{-1}$ | Association | Nucleus | (Shih et al., 2009) |
| **51** | IkBaNFkB => IkBa + NFkB | 6E-5 $min^{-1}$ | Dissociation | Cytoplasm | (Hoffmann et al., 2002) |
| **52** | IkBbNFkB => IkBb + NFkB | 6E-5 $min^{-1}$ | Dissociation | Cytoplasm | (Hoffmann et al., 2002) |
| **53** | IkBeNFkB => IkBe + NFkB | 6E-5 $min^{-1}$ | Dissociation | Cytoplasm | (Hoffmann et al., 2002) |
| **88** | IkBdNFkB => IkBd + NFkB | 6E-5 $min^{-1}$ | Dissociation | Cytoplasm | (Shih et al., 2009) |
| **54** | IkBaNFkBn => IkBan + NFkBn | 6E-5 $min^{-1}$ | Dissociation | Nucleus | (Hoffmann et al., 2002) |
| **55** | IkBbNFkBn => IkBbn + NFkBn | 6E-5 $min^{-1}$ | Dissociation | Nucleus | (Hoffmann et al., 2002) |
| **56** | IkBeNFkBn => IkBen + NFkBn | 6E-5 $min^{-1}$ | Dissociation | Nucleus | (Hoffmann et al., 2002) |
| **89** | IkBdNFkBn => IkBdn + NFkBn | 6E-5 $min^{-1}$ | Dissociation | Nucleus | (Shih et al., 2009) |
| ***IKK-mediated IκB Degradation Reactions*** | | | | | |
| **57** | IkBa => | 0.36 $min^{-1}$ | Prot. Deg. | Cytoplasm | (Mathes et al, 2008) |
| **58** | IkBb => | 0.12 $min^{-1}$ | Prot. Deg. | Cytoplasm | (Mathes et al, 2008) |
| **59** | IkBe => | 0.18 $min^{-1}$ | Prot. Deg. | Cytoplasm | (Mathes et al, 2008) |
| **90** | IkBdIKK1 => IKK1 | 1.2E-3 $min^{-1}$ | Prot. Deg. | Cytoplasm | (Shih et al., 2009) |
| **60** | IkBaNFkB => NFkB | 0.36 $min^{-1}$ | Prot. Deg. | Cytoplasm | (Hoffmann et al., 2002) |
| **61** | IkBbNFkB => NFkB | 0.12 $min^{-1}$ | Prot. Deg. | Cytoplasm | (Hoffmann et al., 2002) |
| **62** | IkBeNFkB => NFkB | 0.18 $min^{-1}$ | Prot. Deg. | Cytoplasm | (Hoffmann et al., 2002) |
| **91** | IkBdIKK1NFkB => IKK1 + NFkB | 0.18 $min^{-1}$ | Prot. Deg. | Cytoplasm | (Shih et al., 2009) |
| ***A20 mRNA and Protein Synthesis and Degradation Reactions*** | | | | | |
| **63** | => A20t (constitutive) | 2 E-6 $min^{-1}$ | RNA Synth. | - | Refer to #1. |
| **64**<br>**65**<br>**66**<br>**71** | => A20t (induced by NFkBn) | 0.4 $\mu M^{-2}$ $min^{-1}$<br>Hill Coefficient: 3.0<br>Delay: 0 min<br>Shutdown: 120 min | RNA Synth. | - | - Refer to #1.<br>- Refer to #1.<br>- Refer to #1.<br>- A20 inducible transcription, as measured by RPA, appears to halt abruptly 2hrs into TNF stimulation. |
| **67** | A20t => | 0.035 $min^{-1}$ | RNA Deg. | Cytoplasm | Refer to #1. |
| **68**<br>**69** | => A20 | 0.25 $min^{-1}$<br>Delay Time: 30 min | Prot. Synth. | Cytoplasm | - Assumed to be equal to IkB translation rates.<br>- Delay was added to account for time between A20 mRNA expression as measured by RPA and A20 protein expression as measured by Western Blot. |
| **70** | A20 => | 0.0029 $min^{-1}$ | Prot. Deg. | Cytoplasm | Supplemental Figure 1 |

| IKK Activation Module | | | | | |
|---|---|---|---|---|---|
| ***TNF-Independent Complex I Activity Reactions*** | | | | | |
| **2** | => tnfrm | 2 E-7 $min^{-1}$ | Prot. Synth. | Cell Surface | Parameter value fit to recapitulate the measured steady-state amount of TNF receptor (Watanabe et al. 1988) |
| **3** | tnfrm => | 0.0058 $min^{-1}$ | Prot. Deg. | Cell Surface | Measured in (Watanabe et al. 1988) |
| **4** | 3 tnfrm => TNFR | 1 E-5 $\mu M^{-1}$ $min^{-1}$ | Association | Cell Surface | Parameter value fit to account for minimal TNF receptor aggregation in the absence of ligand as observed in numerous published studies. |
| **5** | TNFR => 3 tnfrm | 0.1 $min^{-1}$ | Dissociation | Cell Surface | Refer to #4. |
| **6** | TNFR => (internalization) | 0.0017 $min^{-1}$ | Prot. Deg. | Cell Surface | Based upon results published in (Watanabe et al., 1988) showing that the temporal profile of TNF receptor following TNF stimulation. |
| **7** | TNFR + TTR => C1_off | 100 $\mu M^{-1}$ $min^{-1}$ | Association | Cell Surface | Recruitment of TRAF2, TRADD, and RIP adaptors (TTR) to TNFR is required (but not sufficient) for signaling by the TNFR-containing signaling complex (C1). Little biophysical data is available for this reaction; recruitment appears to be simultaneous (Schneider-Brachert et al., 2004). The parameter value represents a compound mechanistic rate constant. It was fit to enable quick activation of downstream IKK activity within the first minutes of stimulation and repression upon removal of TNF ligand in pulse stimulations. |
| **8** | C1_off => TNFR + TTR | 0.75 $min^{-1}$ | Dissociation | Cell Surface | Refer to #7. |
| **9** | C1_off => C1 | 30 $min^{-1}$ | Activation | Cell Surface | The molecular complex containing TNFR, TRAF2, TRADD and RIP undergoes an activation step that involves K63-ubiqutination of RIP. Little biophysical data is available for this step, but parameter fitting was constrained by the fast activation profile of IKK. |
| **10** | C1 => C1_off | 2.0 $min^{-1}$ | Deactivation | Cell Surface | Refer to #9. |
| **11** | C1 => C1_off (A20 mediated) | 1000 $\mu M^{-1}$ $min^{-1}$ | Deactivation | Cell Surface | A20 is known to repress the activity of Complex I. It is a protease of K63-linked ubiquitin chains that deubiquitinates RIP (Wertz et al., 2004). Little biophysical data is available for this step, but parameter fitting was constrained by the IKK activity profiles measured in wild type and $a20^{-/-}$ cells. |
| **12** | C1 => TNFR + TTR | 0.75 $min^{-1}$ | Dissociation | Cell Surface | Assumed to be equal to #8. |
| **13** | C1_off => (internalization) | 0.0017 $min^{-1}$ | Prot. Deg. | Cell Surface | Assumed to be equal to #6. |
| **14** | C1 => (internalization) | 0.0017 $min^{-1}$ | Prot. deg. | Cell Surface | Assumed to be equal to #6. |
| ***TNF-Dependent Complex I Activity Reactions*** | | | | | |
| **1** | tnf => | 0.0154 $min^{-1}$ | Prot. deg. | Extracellular | The half-life of recombinant TNF ligand in cell culture medium was measured by its manufacturer, Roche Diagnostics, to be 45-minutes. |
| **15** | tnf + 3 tnfrm => TNFRtnf | 1100 $\mu M^{-1}$ $min^{-1}$ | Association | Cell Surface | Measured in (Grell et al., 1998). |
| **16** | tnf + TNFR => TNFRtnf | 1100 $\mu M^{-1}$ $min^{-1}$ | Association | Cell Surface | Assumed to be equal to #15. |
| **17** | TNFRtnf => TNFR + tnf | 0.021 $min^{-1}$ | Dissociation | Cell Surface | Measured in (Grell et al., 1998). |
| **18** | TNFRtnf => (internalization | | | | Assumed to be equal to #6. |
| **19** | TNFRtnf + TTR => C1tnf_off | 100 $\mu M^{-1}$ $min^{-1}$ | Association | Cell Surface | Assumed to be equal to #7. TNF binding to the extra-cellular domain of TNFR monomers speeds up trimerization and stabilizes the trimer, but recruitment of the TTR complex to trimerized TNF receptor are assumed to proceed with the same kinetics regardless of the presence of TNF ligand. |
| **20** | C1tnf_off => TNFRtnf + TTR | 0.75 $min^{-1}$ | Dissociation | Cell Surface | Refer to #19. Assumed to be equal to #8. |
| **21** | C1tnf_off => C1tnf | 30 $min^{-1}$ | Activation | Cell Surface | Refer to #19. Assumed to be equal to #9. |
| **22** | C1tnf => C1tnf_off | 2.0 $min^{-1}$ | Deactivation | Cell Surface | Refer to #19. Assumed to be equal to #10. |
| **23** | C1tnf => C1tnf_off (A20 mediated) | 1000 $\mu M^{-1}$ $min^{-1}$ | Deactivation | Cell Surface | Refer to #19. Assumed to be equal to #11. |
| **24** | C1tnf => TNFRtnf + TTR | 0.75 $min^{-1}$ | Dissociation | Cell Surface | Refer to #19. Assumed to be equal to #8. |
| **25** | C1tnf_off => (internalization) | 0.0017 $min^{-1}$ | Prot. deg. | Cell Surface | Refer to #19. Assumed to be equal to #6. |
| **26** | C1tnf => (internalization) | 0.0017 $min^{-1}$ | Prot. deg. | Cell Surface | Refer to #19. Assumed to be equal to #6. |
| **27** | C1tnf_off => C1_off + tnf | 0.021 $min^{-1}$ | Dissociation | Cell Surface | Assumed to be equal to #17. |
| **28** | C1_off + tnf=> C1tnf_off | 1100 $\mu M^{-1}$ $min^{-1}$ | Association | Cell Surface | Assumed to be equal to #15. |
| **29** | C1tnf => C1 + tnf | 0.021 $min^{-1}$ | Dissociation | Cell Surface | Assumed to be equal to #17. |
| **30** | C1 + tnf => C1tnf | 1100 $\mu M^{-1}$ $min^{-1}$ | Association | Cell Surface | Assumed to be equal to #15. |

| | | | | | |
|---|---|---|---|---|---|
| ***IKKK (TAB1/2-TAK1 complex) Activity Reactions*** | | | | | |
| **31** | IKKK_off => IKKK (constitutive) | 5 E-7 min$^{-1}$ | Activation | Cytoplasm | Parameter value fit to account for low IKK activity in the absence of ligand as measured by IKK Kinase Assay (O'Dea et al., 2007). |
| **32** | IKKK_off => IKKK (C1 mediated) | 500 μM$^{-1}$ min$^{-1}$ | Activation | Cytoplasm | Refer to #7. |
| **33** | IKKK_off => IKKK (C1tnf mediated) | 500 μM$^{-1}$ min$^{-1}$ | Activation | Cytoplasm | Refer to #19. Assumed to be equal to #32. |
| **34** | IKKK => IKKK_off (constitutive) | 0.25 min$^{-1}$ | Deactivation | Cytoplasm | The constitutive inactivation rate of this complex was fit to ensure low basal activity and efficient repression following TNF pulse stimulation. |
| ***IKK Activity Reactions*** | | | | | |
| **35** | IKK_off => IKK | 5 E-5 min$^{-1}$ | Activation | Cytoplasm | Refer to #31 |
| **36** | IKK_off => IKK (IKKK mediated) | 520 μM$^{-1}$ min$^{-1}$ | Activation | Cytoplasm | Refer to #7. |
| **37** | IKK => IKK_off | 0.02 min$^{-1}$ | Deactivation | Cytoplasm | Refer to #34. |
| **38** | IKK => IKK_i (self-inactivation) | 0.15 min$^{-1}$ | Deactivation | Cytoplasm | IKK is thought to down-regulate its own activity via auto-phosphorylation of C-terminal serine residues (Delhase et al., 1999). This mechanism was not shown to cause IKK protein degradation and is distinct from inactivating IKK via constitutive phosphatase activity (Refer to #94). The parameter value was fit to temporal profiles of IKK activity in response to TNF stimulation (Werner et al., 2005). |
| **39** | IKK_i => IKK_off | 0.02 min$^{-1}$ | Deactivation | Cytoplasm | C-terminally phosphorylated IKK is assumed to be subject to constitutive phosphatase activity. Refer to #38. |

| | | | | | |
|---|---|---|---|---|---|
| **Necroptosis module** | | | | | |
| ***Necroptosis Activity Reactions*** | | | | | |
| **92** | C1_off + C1 --> RIP1 | 6 min$^{-1}$ | Activation | Cytoplasm | Fitted to data of RIP1 protein |
| **93** | A20 --\| RIP3 | 6 min$^{-1}$ | Inhibition | Cytoplasm | Fitted to data of A20 protein and RIP3 protein |
| **94** | RIP1 => | 0.6 μM$^{-1}$ min$^{-1}$ | Degradation | Cytoplasm | Fitted to data of RIP1 protein |
| **95** | RIP1 --> RIP3 | 6 min$^{-1}$ | Activation | Cytoplasm | Fitted to data of RIP1 protein and RIP3 protein |
| **96** | RIP3 => | 6 μM$^{-1}$ min$^{-1}$ | Degradation | Cytoplasm | Fitted to data of RIP3 protein |
| **97** | RIP3 --> pMLKL | 300 min$^{-1}$ | Activation | Cytoplasm | Fitted to data of RIP3 protein and pMLKL protein |
| **98** | pMLKL => | 60 μM$^{-1}$ min$^{-1}$ | Degradation | Cytoplasm | Fitted to data of pMLKL protein |

Note for Supplementary Table S6: There are 98 equations in the model. The NF$\kappa$B module and IKK activation module are modeled by chemical reactions by the law of mass action. The necroptosis module is coarse-grained as activation and inhibition regulatory network, which are modeled by using Hill functions.

#### Supplementary Table S7: Perturbed parameters for NF$\kappa$B module

| # in NF$\kappa$B model | Parameter | Perturbed ratio | Adjustment on Parameter Value |
|---|---|---|---|
| **1** | TNF decay rate | 0.1 | Fitted to death time distribution |
| **63** | A20 basal transcription | 0.6 | Fitted to data of A20 mRNA for L929 cells |
| **64** | A20 induced transcription | 0.15 | Fitted to data of A20 mRNA for L929 cells |
| **68** | A20 translation | 20 | Fitted to data of A20 protein for L929 cells |
| **2** | I$\kappa$B$\beta$ basal transcription | 0.85 | Fitted to data of I$\kappa$B$\beta$ mRNA for L929 cells |
| **5** | I$\kappa$B$\beta$ induced transcription | 20 | Fitted to data of I$\kappa$B$\beta$ mRNA for L929 cells |
| **72** | I$\kappa$B$\delta$ basal transcription | 0.3 | Fitted to data of I$\kappa$B$\delta$ mRNA for L929 cells |
| **73** | I$\kappa$B$\delta$ induced transcription | 4 | Fitted to data of I$\kappa$B$\delta$ mRNA for L929 cells |

#### Supplementary Table S8: Parameters for necroptosis module

| Reaction # | Parameter | Parameter Value | Adjustment on Parameter Value |
|---|---|---|---|
| **92** | Synthesis rate $k_{s1}$ | distributed | |
| **95** | Synthesis rate $k_{s2}$ | 6 min$^{-1}$ | Fitted to data of RIP3 protein |
| **97** | Synthesis rate $k_{s3}$ | 300 min$^{-1}$ | Fitted to data of RIP3 protein and pMLKL protein |
| **94** | Degradation rate $k_{d1}$ | 0.6 μM$^{-1}$ min$^{-1}$ | Fitted to data of RIP3 protein |
| **96** | Degradation rate $k_{d2}$ | 6 μM$^{-1}$ min$^{-1}$ | Fitted to data of RIP3 protein |
| **98** | Degradation rate $k_{d3}$ | 60 μM$^{-1}$ min$^{-1}$ | Fitted to data of pMLKL protein |
| **92** | $Km_1$ | 1 μM | Fitted to data of RIP3 protein |
| **95** | $Km_2$ | 0.15 μM | Fitted to data of RIP3 protein |
| **93** | $Km_3$ | 0.5 μM | Fitted to data of A20 protein and RIP3 protein |
| **97** | $Km_4$ | 0.3 μM | Fitted to data of RIP3 protein and pMLKL protein |
| **92** | Hill coefficient $n_1$ | 3 | Fitted to data of RIP3 protein |
| **95** | Hill coefficient $n_2$ | 3 | Fitted to data of RIP3 protein |
| **93** | Hill coefficient $n_3$ | 5 | Fitted to data of A20 protein and RIP3 protein |
| **97** | Hill coefficient $n_4$ | 3 | Fitted to data of RIP3 protein and pMLKL protein |

#### Supplementary Table S9: Distributed parameters for necroptosis model

| Reaction # | Parameter | Parameter distribution |
|---|---|---|
| **92** | Synthesis rate $k_{s1}$ from RIP1 to RIP3 | lognrnd(0,0.5)*0.7, where lognrnd(a,b) is log normal distribution with mean a and variable b. |
| **64** | A20 induced transcription | 0.0053*normal(3,30)+ 0.0036* normal(600,135) , where normal(a,b) is a normal distribution with mean a and variable b. |
